# The tree that hides the forest: identification of common predisposing loci in several hematopoietic cancers and several dog breeds

**DOI:** 10.1101/2020.07.23.214007

**Authors:** Benoit Hédan, Edouard Cadieu, Maud Rimbault, Amaury Vaysse, Patrick Devauchelle, Nadine Botherel, Jérôme Abadie, Pascale Quignon, Thomas Derrien, Catherine André

## Abstract

Histiocytic sarcoma (HS) is a rare but aggressive cancer in humans and dogs. The spontaneous canine model, with the clinical, epidemiological and histological similarities with human HS and specific breed predispositions, is a unique model/opportunity to unravel the genetic bases of this cancer. In this study, we aimed to identify germline risk factors associated with the development of HS in canine predisposed breeds. We used a methodology that combined several genome-wide association studies in a multi-breed and multi-cancer approach, as well as targeted next generation sequencing, and imputation combining several breeds (Bernese mountain dog, Rottweiler, flat coated retriever and golden retriever) and three haematopoietic cancers (HS, lymphoma and mast cell tumor). Results showed that we not only refined the previously identified HS risk *CDKN2A* locus but we identified new loci on canine chromosomes 2, 5, 12, 14, 20, 26 and X. Capture and targeted sequencing of specific loci pointed towards the existence of regulatory variants in non coding regions and/or methylation mechanisms linked to risk haplotypes, leading to strong cancer predispositions in specific dog breeds. Our results showed that these canine cancer predisposing loci appear to be due to the additive effect of several risk haplotype involved also in other haematopoietic cancers such lymphoma or mast cell tumor, illustrating the pleiotropic nature of these canine cancer loci as observed in human oncology, thus reinforcing the interest of predisposed dog breeds to study cancer initiation and progression.

## 1. Introduction

Over the past decade, the Dog has arisen as a relevant and under used spontaneous model for the analysis of cancer predisposition and progression, as well as development and trials of more efficient therapies for a number of human cancers^1–10^. With over 4.2 million dogs diagnosed with cancer annually in USA^8^, canine cancers represent a unique source of spontaneous tumors. Canine cancers share strong similarities with human’s tumors, both on biological behavior and histopathological features^11–14^. Spontaneous canine models thus appear a natural and ethical *i.e*. non-experimental model to decipher the genetic bases of cancers. Indeed, in most human cancers, presenting incomplete penetrance and genetic heterogeneity, identifying the genetic predisposition is complex^6^, and almost impossible for rare cancers. With their specific breed structures and artificial selection, dog breeds have gained numerous susceptibilities to genetic diseases and a limited number of critical genes are involved in complex diseases, and even in cancers^6^. Numerous genome wide association studies (GWAS) in dogs illustrate that in complex traits, such as body size or cancer, a small number of loci with strong effect are involved in dogs, as compared to humans, facilitating their identification. With large intra-breed linkage disequilibrium LD, cancer loci were successfully identified even with a small number of cases and controls^15–20^. Thus, spontaneously affected pet dogs, with breed specific cancers, appear as efficient natural models to identify the genetics underpinning several dog-human homologous cancers.

In humans, histiocytic sarcoma (HS) is extremely rare, involving histiocytic cells (dendritic or monocytic/macrophagic lineages) with a limited response to chemotherapy and a high mortality. Due to the rarity of this cancer, there is no consensus on prognostic factors and standard treatment^21^ and models are strongly needed to better understand this dramatic cancer. In the whole dog species, HS is a relatively rare cancer, but a few popular breeds are highly predisposed to this cancer: Bernese Mountain Dogs (BMD), Rottweiler, Retrievers (especially Flat Coated Retrievers –FCR-) with specific clinical presentations per breed. The clinical presentation, histopathology and issue of this canine cancer are similar to those observed in humans^22,23^. These breed predispositions allowed the collection of numerous samples to successfully identify somatic variants of the MAPK pathway^24,25^ and we recently showed that the same mutations of *PTPN11*, the most frequently altered gene of the MAPK pathway, are found in human and canine HS^25^. But most importantly, these breed predispositions to specific clinical presentations allow to unravel the genetic bases of HS. Indeed, we previously showed that the *PTPN11* somatic mutations found in half of HS canine cases are linked to a HS aggressive clinical subgroup in dogs and in humans^25^. Regarding predisposition to HS, previous work on 236 cases and 228 controls allowed to highlight the *MTAP-CDKN2A* genomic region as one of the main locus conferring susceptibility to HS in BMD^26^. Nevertheless, HS is a multifactorial cancer, and other loci are expected to be involved in HS predisposition. This is in accordance with secondary hits already observed in the previous GWAS analyses^26^. Furthermore, despite a strong/important heritability of HS in BMD^27^, the awareness of this devastating cancer and attempt to selection against HS since 20 years, breeders have not succeeded to reduce the prevalence of this cancer. In addition, it is suspected that HS predisposed breeds (BMD, Rottweiler and retrievers) share common risk alleles due to common ancestors, thus cases from close breeds can accelerate the identification of common loci by reducing the haplotype of these critical regions^28^. Interestingly, these HS predisposed breeds also present a high risk of developing other cancers: it is estimated that a high proportion of deaths in BMD (45-76 %), Golden retriever (39-50%), Labrador retriever (31-34%), Flat Coated Retriever (54%) and Rottweiler (30-45%) are due to several neoplasms^29–31^. Indeed, these breeds are also prone to develop lymphoma, mast cell tumors, hemangiosarcoma, osteosarcoma, or melanoma^26,31,32^.

This study aims to extend previous studies to decipher the genetic bases of HS, based on a multi-breed approach. We thus performed exhaustive GWAS with increased numbers of cases and controls from three different breeds and with higher density SNV arrays. These data led us to identify important loci, complementary to the *MTAP-CDKN2A* locus, and showed that they are also associated to other cancers in predisposed breeds. Our study not only confirms the main role of the *CDKN2A* locus in HS predisposition by identifying shared variants between the predisposed breeds but also sheds light on secondary loci located on canine chromosomes 2, 5, 12, 14, 20, 26 and X containing relevant candidate genes. The effect of these loci appears to be due to the additive effect of sub-loci involved also in other haematopoietic cancers such lymphoma or mast cell tumor. Capture and targeted sequencing of specific loci did not allow the identification of straightforward variants linked to cancer predispositions. Finally, our results point towards the existence of regulatory variants in non coding regions and/or methylation mechanisms linked to risk haplotypes, leading to strong cancer predispositions in specific dog breeds.

## 2. Results

To decipher the genetic bases of histiocytic sarcoma (HS) in the dog model system, we took advantage of the well known HS predisposed breeds. To this aim, we combined data from GWAS with high-density genotyped and imputed SNV data in the following breeds: BMD, FCR, Rottweiler with the following cancers: HS, lymphoma and mast cell tumor, as well as public available data from lymphoma and mast cell tumor in golden retriever^15,16^ (Table 1).

**Table 1:** Characteristics of the GWAS analyses performed in this study

| GWAS Name | Paragrah | cancers | breeds | Number of cases | Number of controls | SNP Arrays used |
| --- | --- | --- | --- | --- | --- | --- |
| GWAS_1_HS_BMD | §3.1 | HS | BMD | 172 | 154 | Illumina 173k SNV Canine HD array |
| GWAS_2_HS+lymphoma_BMD | §3.2a | HS and lymphoma | BMD | 252 | 146 | Illumina 173k SNV Canine HD array |
| GWAS_3_HS+lymphoma_BMD+golden_retriever | §3.2a | HS and lymphoma | BMD, golden retriever | 293 | 318 | Illumina 173k SNV Canine HD array |
| GWAS_4_HS+MCT_BMD | §3.2b | HS and mast cell tumor | BMD | 216 | 135 | Illumina 173k SNV Canine HD array |
| GWAS_5_HS+MCT_BMD+golden_retriever | §3.2b | HS and mast cell tumor | BMD, golden retriever | 281 | 197 | Illumina 173k SNV Canine HD array |
| GWAS_6_HS_BMD+FCR | §3.3 | HS | BMD, FCR | 185 | 169 | Illumina 173k SNV Canine HD array |
| GWAS_7_HS_BMD+Rottweiler | §3.3 | HS | BMD, Rottweiler | 210 | 176 | Illumina 173k SNV Canine HD array |
| GWAS_8_HS_BMD_with_imputed_SNV | §3.4 | HS | BMD | 404 | 371 | Affymetrix Axiom Canine Genotyping Array 1.1M SV<br>Illumina 20k SNV Canine HD array imputed for 1.1M SV<br>Illumina 173k SNV Canine HD array imputed for 1.1M SV |
| GWAS_9_HS_BMD+FCR_with_imputed_SNV | §3.4 | HS | BMD, FCR | 417 | 386 | Affymetrix Axiom Canine Genotyping Array 1.1M SV<br>Illumina 20k SNV Canine HD array imputed for 1.1M SV<br>Illumina 173k SNV Canine HD array imputed for 1.1M SV |
| GWAS_10_HS_BMD+FCR+Rottweiler_with_imputed_SNV | §3.4 | HS | BMD, FCR,Rottweiler | 455 | 408 | Affymetrix Axiom Canine Genotyping Array 1.1M SV<br>Illumina 20k SNV Canine HD array imputed for 1.1M SV<br>Illumina 173k SNV Canine HD array imputed for 1.1M SV |

### 2.1. Identification of loci linked to the risk to develop HS in the BMD breed

Using Bernese Mountain Dog DNA from 172 HS cases and 154 controls (older than 10 years), we performed a first round of genome-wide association study (GWAS), correcting for population stratification and cryptic relatedness. We thus identified 29 SNVs significantly associated to HS with one SNV localized on chromosome 5 (CFA5:30496048, *p*_corrected_ = 6.36 x 10^−5^), 27 SNVs on chromosome 11 (CFA11:41161441, *p*_corrected_ = 4.85 x 10^−7^) and one SNV on chromosome 20 (CFA20:30922308, *p*_corrected_ = 2.52 x 10^−5^). This GWAS confirmed that the main locus linked to HS is located on CFA11, overlapping the *MTAP-CDKN2A* region, a locus previously associated with HS^26^. The present analysis also identified new loci on CFA5 and CFA20 (Figure 1). Interestingly, these three regions were previously identified in the predisposition of other cancers in dogs: CFA11 in osteosarcoma, CFA5 in lymphoma and hemangiosarcoma, CFA20 in mast cell tumor^15–17^. Indeed Tonomura et al identified two independent peaks on CFA5, involved in lymphoma and hemangiosarcoma^16^, overlapping the CFA5 locus found in HS; while Arendt et al identified at least two independent peaks on CFA20 involved in mast cell predisposition^15^, also overlapping the CFA20 locus found in HS of the Bernese Mountain dog breed. Thus, we hypothesized that, due to the strong breed selection, the significant associations detected for HS could be due to cumulative risk alleles/haplotypes than can also be at risk for other haematopoietic cancers. This hypothesis is reinforced by the fact that BMDs also show a significant predisposition to lymphoma^33^ and that in HS affected BMDs families, we observed the segregation of other hematopoietic cancers such as mast cell tumor and lymphomas, with high frequency (Figure 2).

**Figure 1:**
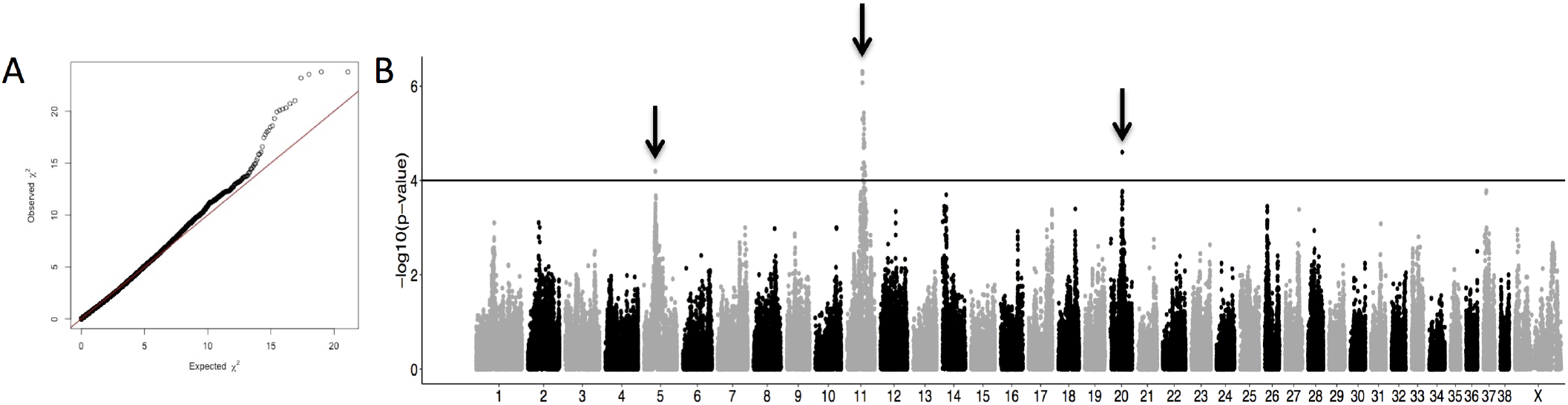
Results of Genome-wide association studies (GWAS) on BMD with 172 HS cases and 154 controls (GWAS_1_HS_BMD). A) Quantile-Quantile plot displaying a genomic inflation λ of 1.000005, indicating no residual inflation. B) Manhattan plot displaying the statistical results from the GWAS. This analysis pointed out three loci (arrows) on chromosome 5 (CFA5:30496048, *p*_corrected_ = 6.36 x 10^−5^), on chromosome 11 (CFA11:41161441, *p*_corrected_ = 4.85 x 10^−7^) and on chromosome 20 (CFA20:30922308, *p*_corrected_ = 2.52 x 10^−5^).

**Figure 2:**
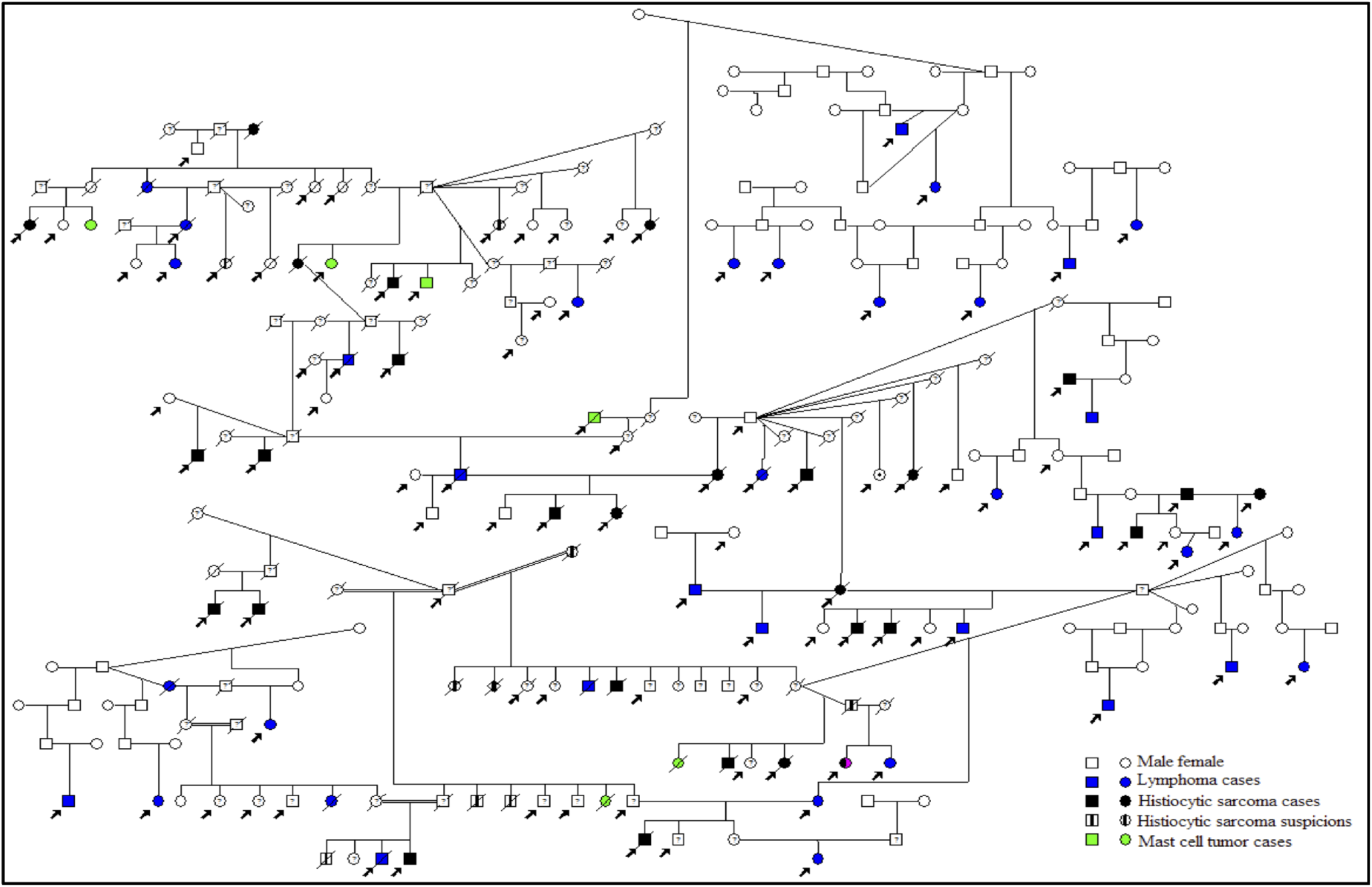
Pedigrees of a Bernese Mountain dog family showing the co-segregation of lymphoma (blue) and mast cell tumor (green) with histiocytic sarcoma (black).

### 2.2. Involvment of HS loci in other haematopoietic cancers

To test if these HS predisposing loci could be also involved in the predisposition of lymphoma or mast cell tumor, we added lymphoma or mast cell tumor cases to the previous HS GWAS.

#### a. Lymphoma

The addition of 80 lymphoma affected BMDs to the first GWAS (GWAS_1_HS_BMD) identified 14 significantly associated SNVs, of which 5 were located on the CFA5 between 30.2Mb and 32.2Mb with the most significant SNV being at 30.5 Mb (best SNV CFA5: 30496048, *p*_corrected_ = 3.13×10^−6^) (Figure 3 and Supplementary Figure 1). This result confirmed that the CFA5 locus is common to HS and lymphoma predisposition in BMD. The top SNV of CFA5 locus is located in an intron of the *SPNS3* gene (sphingolipid transporter 3) for which the paralogous gene (*SPNS2*) is known to be important in immunological development, playing a critical role in inflammatory and autoimmune diseases (https://www.ncbi.nlm.nih.gov/gene/124976). Since this locus overlapped the two lymphoma predisposing peaks (29.8 Mb and 33 Mb respectively) found by Tonomura *et al* in the Golden retriever, we performed a meta-analysis, including BMD GWAS data from this study and the publicly available golden retriever GWAS data^16^. The addition of golden retriever lymphoma cases (n=41) and controls (n=172) to the second BMD GWAS (GWAS_2_HS+lymphoma_BMD), containing 252 HS or lymphoma cases and 146 controls, resulted in an increased signal on the CFA5 locus, strengthening the two loci previously identified by Tonomura *et al* at 29.8 Mb and 33 Mb (SNV CFA5: 29836124, *p*_corrected_ =4.75×10^−7^ and SNV CFA5: 33001550, *p*_corrected_ =8.96 x10^−8^) (Figure 3 and Supplementary Figure 1). These results show that BMDs and golden retrievers share common risk loci on CFA5 involved in hematopoietic cancers and that the CFA5 association with HS and lymphoma in BMD is due to additional effects of at least 3 different peaks on CFA 5 (30.5 Mb, 29.8 Mb and 33 Mb) (Figure 3).

**Figure 3:**
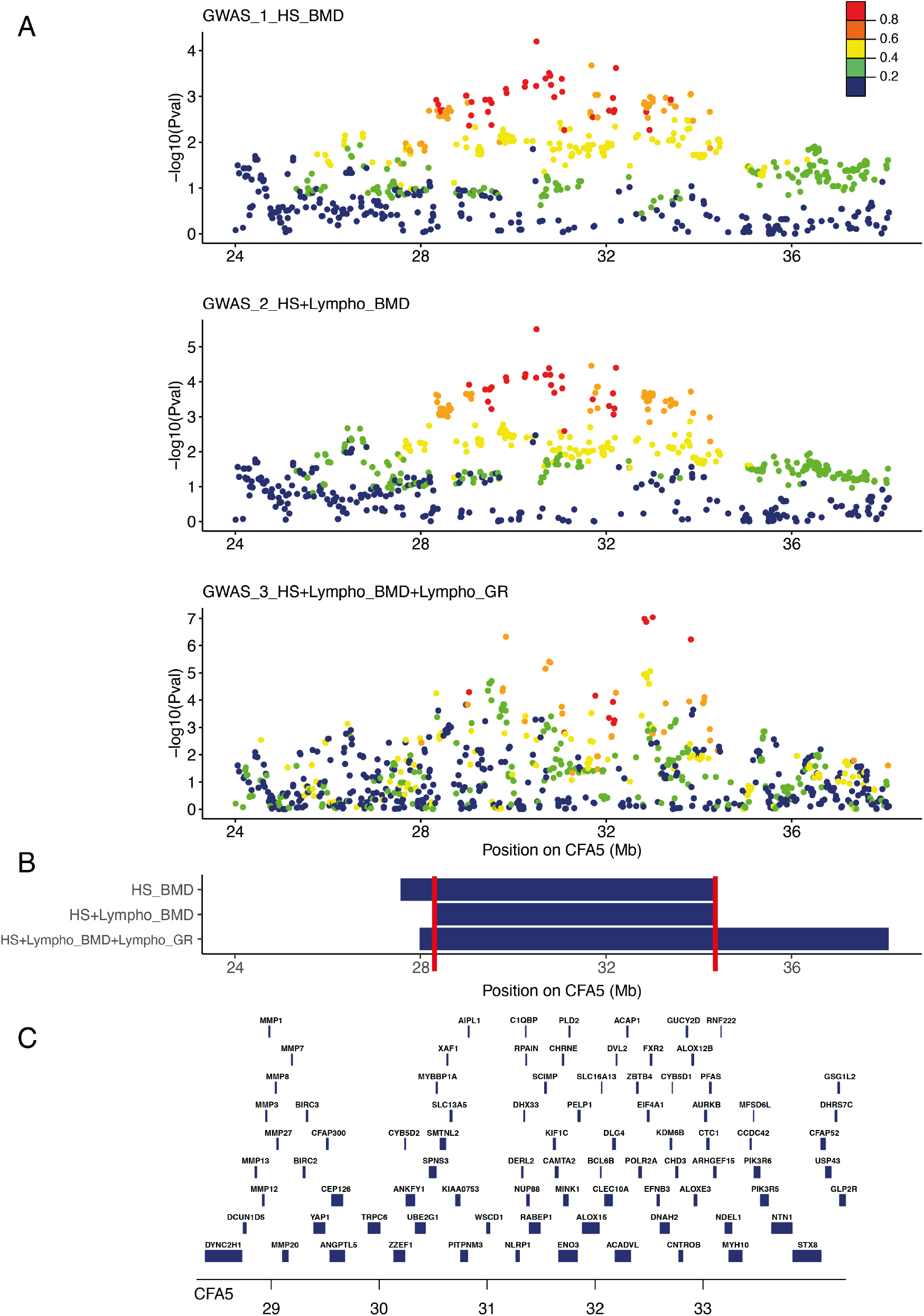
Close up view of the CFA5 locus: A. Manhattan plot on the CFA5 20-40 Mb region highlighting the best p-values obtained in the 3 GWAS experiments and the R^2^ in cases of the SNVs with the top GWAS SNV are depicted to show LD structure: BMD GWAS for HS with 172 cases vs 154 controls (GWAS_1_HS_BMD); BMD GWAS for HS and lymphoma with 252 cases vs 146 controls (GWAS_2_HS+lymphoma_BMD, black dots); meta-analysis combining the BMD GWAS for HS and lymphoma (252 cases vs 146 controls) and the golden retriever GWAS for lymphoma (41 cases vs 172 controls) from Tonomura *et al*. 2015 (GWAS_3_HS+lymphoma_BMD+golden_retriever, green dots). B. Regions delimitated by SNPs in LD with the best GWAS SNPs (R^2^ >0.6) in cases, the minimal region between the three GWAS (CFA5: 28309815-34321500) is delimitated by red lines. C. Close up view of the genes (with available symbol) located in this minimal 28-34Mb region of CFA5.

#### b. Mast cell tumor

The addition of 44 BMDs with mast cell tumor (MCT) to the first HS BMD GWAS (GWAS_1_HS_BMD) identified 11 significantly associated SNVs (best SNV CFA11: 41161441, *p*_corrected_ = 1.31 x10^−6^) of which 4 were located on the CFA20 (best SNV on CFA20 CFA20: 30922308, *p*_corrected_ = 4.05 x10^−6^) (Figure 4 and Supplementary Figure 2). This result confirmed that the CFA20 locus is common to HS and MCT predisposition in BMD. The top SNV of CFA20 locus lies in an intron of *FHIT*, a tumor suppressor involved in apoptosis and prevention of the epithelial-mesenchymal transition^34^. Interestingly, this locus overlapped one of the three MCT independent predisposing peaks (33Mb, 39Mb and 45Mb Canfam3) identified in the golden retriever^15^. We then performed a meta-analysis combining our BMD GWAS for HS and MCT (GWAS_4_HS+MCT_BMD) with the golden retriever GWAS for MCT, by adding the publicly available data of Arendt et al. 2015. The addition of European golden retriever mast cell tumor cases and controls resulted in an increase association signal in the CFA20 locus, clearly pointing out the CFA20 locus, at 33Mb (2 best SNVs: CFA20: 33321282, *p*_corrected_ =3,42 x10^−8^, CFA20: 31940114, *p*_corrected_ =2,34 x10^−7^ in *ARHGEF3* genes and close to *IL17RD* gene), one of the three peaks identified by Arendt et al. 2015. These results show that the BMDs and golden retriever breeds share common inherited risk factors on CFA20 for histiocytic sarcoma and for MCT cancers and that the CFA20 association with cancers in BMD is due to additional effects of at least two peaks (Supplementary Figure 2 and Figure 4). These results also confirmed that CFA20 association with MCT in golden retriever is due to the additional effect of at least 3 risk haplotypes (33Mb, 39Mb and 45 Mb).

**Figure 4.**
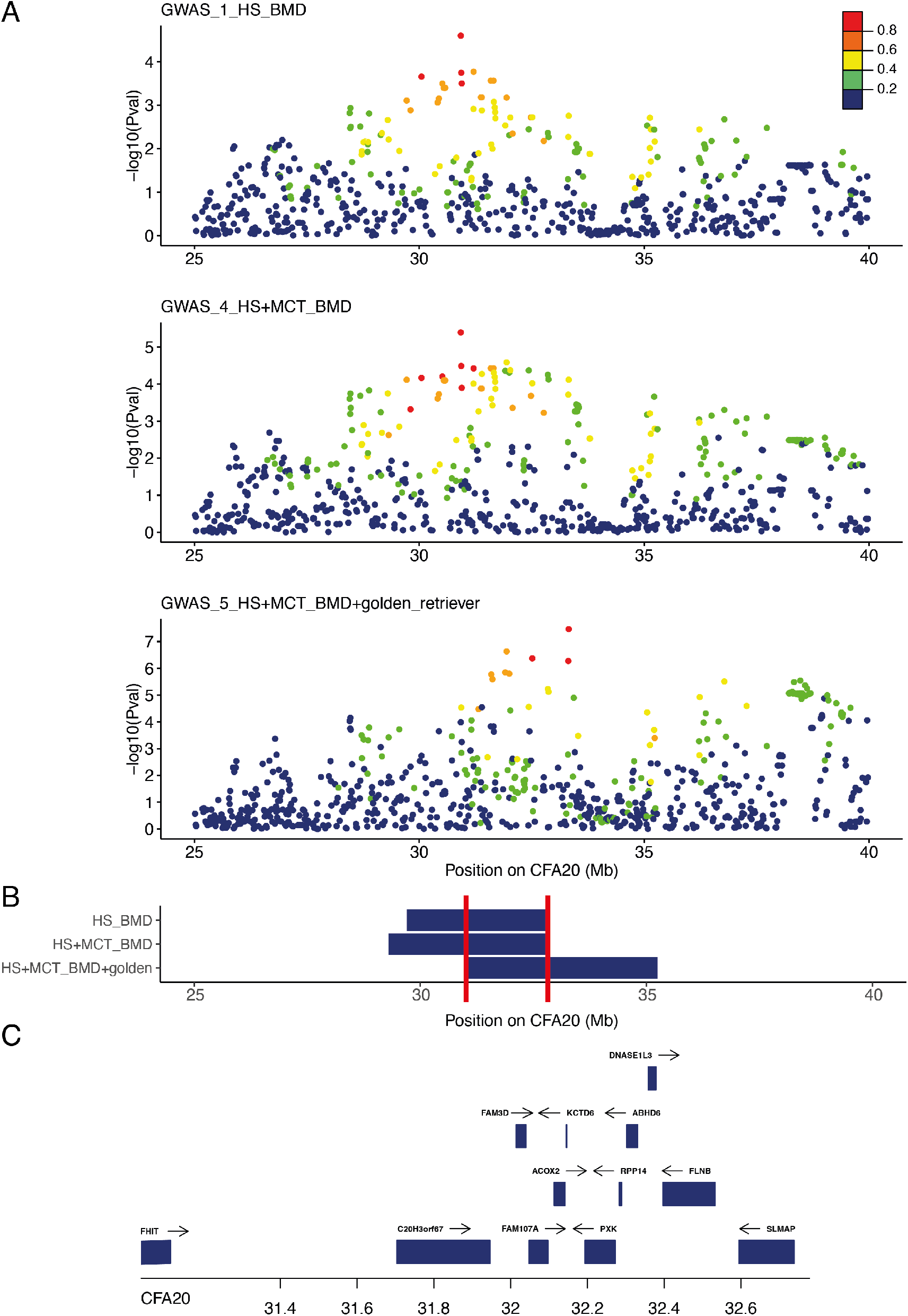
Close up view of the CFA20 locus: A. Manhattan plot on the CFA20 20-45 Mb region highlighting the best p-values obtained in the 3 GWAS experiments and the R^2^ in cases of the SNVs with the top GWAS SNV are depicted to show LD structure: BMD GWAS for HS with 172 cases vs 154 controls (GWAS_1_HS_BMD); BMD GWAS, for HS and mast cell tumor with 216 cases vs 135 controls (GWAS_4_HS+MCT_BMD); the meta-analysis combining BMD GWAS for HS and mast cell tumor with the European golden retriever (65 cases vs 62 controls) from Arendt et *al*. 2015 (GWAS_5_HS+MCT_BMD+golden_retriever). B. Regions delimitated by SNPs in LD with the best GWAS SNPs (R^2^ >0.6) in cases, the minimal region between the three GWAS (CFA20: 31036863-32778949) is delimitated by red lines. C. Close up view of the genes (with available symbol) located in this 31-33Mb minimal region of CFA20.

To conclude, it appears that the loci linked to cancer in dogs could have pleiotropic effects-associated with the risk of several cancers in several breeds- and be due to the additive effects of several risk peaks. Thus, to identify more precisely HS risk haplotypes and to reduce the loci size, we added to the first BMD GWAS for HS (GWAS_1_HS_BMD) new HS cases from other HS predisposed breeds.

### 2.3. Refining HS Loci by multiple-breed analyses

The addition of 28 Flat Coated Retrievers (FCR) (13 HS cases and 15 controls) to the first BMD GWAS for HS (GWAS_1_HS_BMD) identified 22 significantly associated SNVs located on the CFA11 (best SNV CFA11: 44796340, *p*_corrected_ = 1.6 x10^−6^). The reduced signal observed on CFA11 at 41 Mb with the addition of the FCR compared to BMD alone, was not in favor of a shared risk haplotype between FCR and BMD on this locus (Figure 5B).

**Figure 5:**
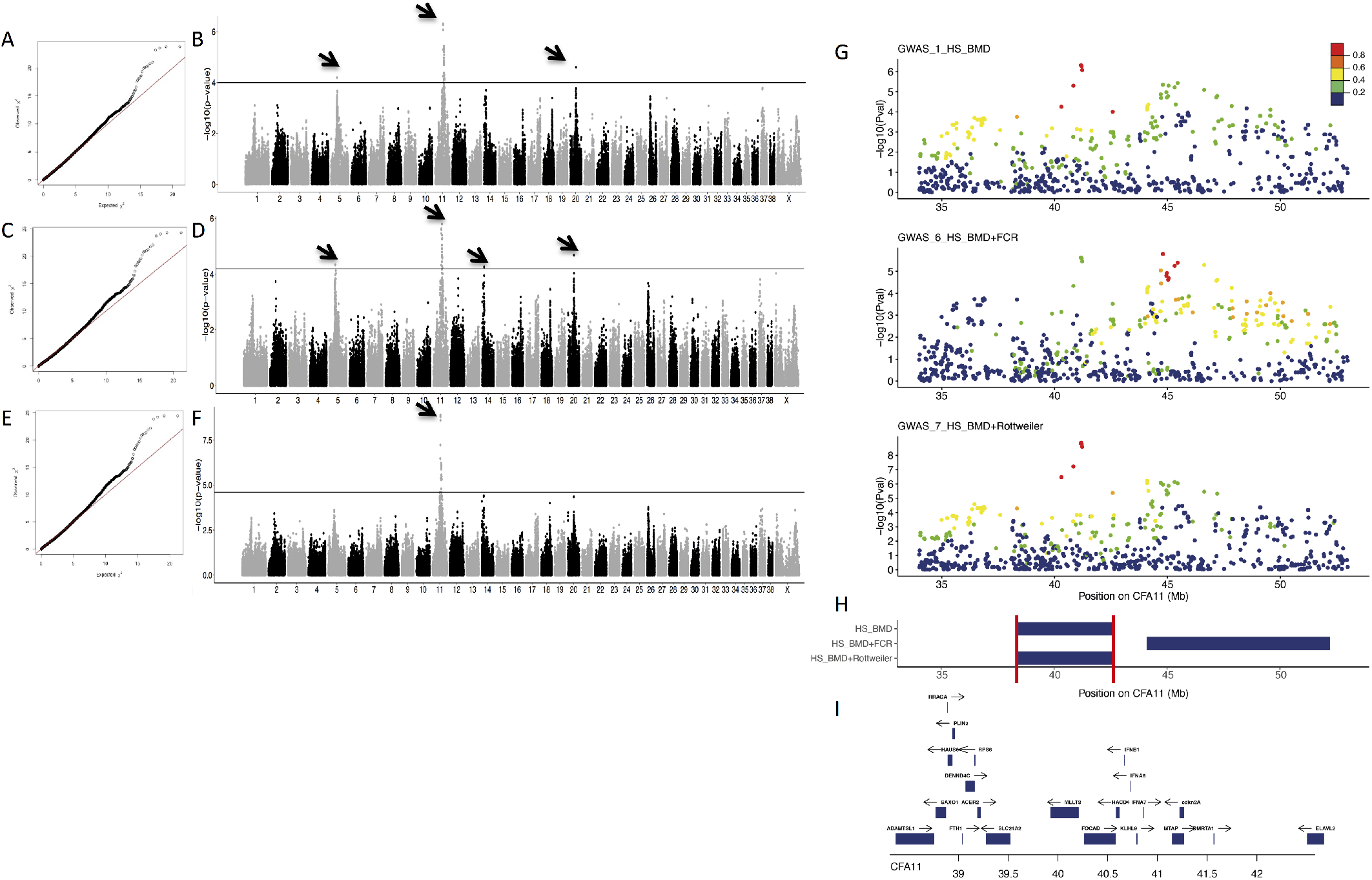
Genome-wide association studies on BMD and other predisposed breeds on HS. A & B: BMD GWAS results for HS with 172 cases and 154 controls (GWAS_1_HS_BMD). A) Quantile-Quantile plot displaying a genomic inflation λ of 1.000005, indicating no residual inflation. B) Manhattan plot displaying the statistical results from the GWAS. This analysis pointed out three loci (arrows) on chromosome 5 (CFA5: 30496048, *p*_corrected_ = 6.36 x 10^−5^), on chromosome 11 (CFA11: 41161441, *p*_corrected_ = 4.85 x 10^−7^) and on chromosome 20 (CFA20:30922308, *p*_corrected_ = 2.52 x 10^−5^). C & D: GWAS results for HS combining BMDs (172 cases vs 154 controls) and FCRs (13 cases vs 15 controls) (GWAS_6_HS_BMD+FCR). C) Quantile-Quantile plot displaying a genomic inflation λ of 1.000004, indicating no residual inflation. D) Manhattan plot displaying the statistical results from the GWAS. This analysis pointed out the loci on four loci (arrows) on chromosome 5 (CFA5: 30496048, *p*_corrected_ = 4.64 x 10^−5^), chromosome 11 (CFA11: 44796340, *p*_corrected_ = 1.6 x 10^−6^), chromosome 14 (CFA14: 10334651, *p*_corrected_ = 5.55 x 10^−5^) and chromosome 20 (CFA20: 30922308, *p*_corrected_ = 2.1 x 10^−5^). E & F: GWAS results for HS combining BMDs (172 cases vs 154 controls) and Rottweilers (38 cases vs 22 controls) (GWAS_7_HS_BMD+Rottweiler). E) Quantile-Quantile plot displaying a genomic inflation λ of 1.000004, indicating no residual inflation. F) Manhattan plot displaying the statistical results from the GWAS. This analysis pointed out the locus on chromosome 11 (arrow) (CFA11: 41176819, *p*_corrected_ = 1.28 x 10^−9^). G) Close up view of the CFA11 locus highlighting the best p-values obtained in the three GWAS experiments and the R^2^ in cases of the SNVs with the top GWAS SNV are depicted to show LD structure: BMDs GWAS (GWAS_1_HS_BMD), BMDs plus FCRs GWAS (GWAS_6_HS_BMD+FCR), BMDs plus Rottweilers GWAS (GWAS_7_HS_BMD+Rottweiler). H) Regions delimitated by SNPs in LD with the best GWAS SNPs (R^2^ >0.6) in cases, the minimal region between the three GWAS (CFA11: 38308478-42582141) is delimitated by red lines. I) Close up view of the genes (with available symbol) located in this 38-42 Mb minimal region.

The addition of 60 Rottweilers (38 cases and 22 controls) to the first BMD GWAS for HS (GWAS_1_HS_BMD) identified 21 significantly associated SNVs located on CFA11 (best SNV CFA11: 41176819, *p*_corrected_ = 1.28 x10-9) (Figure 5D). The increased signal observed on CFA11 at 41 Mb confirmed that the CFA11 locus at 41Mb is common to BMDs and Rottweilers.

Haplotype analysis confirmed that BMD and Rottweilers share a common at risk haplotype tagged by 7 SNVs spanning 2.3Mb (CFA11: 40298068-42570967) while the analysis of FCRs pointed out a different risk haplotype (Table 2). This BMD-Rottweiler shared haplotype appears specific to BMD and Rottweilers HS cases since it is never found in other 231 Swiss dog breeds (18 Appenzeller Sennenhund, 8 Entlebucher Mountain dog and 205 greater Swiss dog, data not shown), breeds phylogenetically closed to BMD but not affected by HS.

**Table 2:**
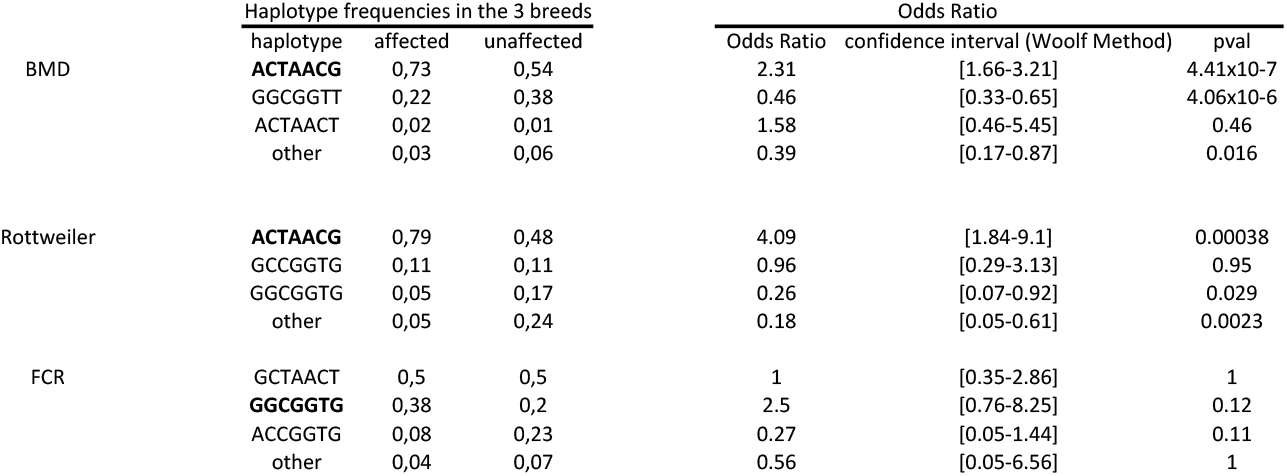
Association analysis between the haplotypes and the phenotypes in predisposed breeds. The haplotype was determined with the genotypes of seven CFA11 SNVs located at 40298068, 40831903, 41161441, 41176819, 41196587, 41217026, 42570967 bp. At risk haplotypes in the breed are represented in bold. CI: confidence interval (Woolf Method). BMD: Bernese Mountain Dog; FCR: Flat Coated Retriever.

### 2.4. Refining HS Loci by imputation on higher density SNV array

The fact that no shared risk haplotype was found between FCR and other HS predisposed breeds on the CFA11 *MTAP-CDKN2a* locus (at 41 Mb) could be due to a lack of density of the Illumina 173K SNV Canine HD array, resulting in the impossibility to identify a short risk haplotype. To test this hypothesis, 134 dogs were genotyped on the high density Affymetrix Axiome Canine Genotyping array (1.1M SNV) and all SNVs were imputed on the Illumina 173K SNV Canine HD. In addition, to increase the power of the GWAS, we added previously published BMD cases and controls^26^ genotyped on the Canine SNP20 Bead-Chip panel (Illumina −22k SNV) and also imputed the genotypes on the high density Axiome Canine Genotyping array (1.1M SNV).

The addition of BMD cases and controls to the first BMD GWAS (GWAS_1_HS_BMD), resulted in a total of 404 cases and 371 controls imputed on 592,073 SNVs. The statistical analysis allowed the identification of 2,103 SNVs significantly associated to HS (Table 3, Figure 6 AB). This GWAS, by increasing the number of BMD cases and controls, confirmed the involvement of the CFA11 locus as well as the role of other loci (CFA5 and CFA14) in HS BMD predisposition and identified new loci on chromosomes 2, 12, 26 and X, close to relevant candidates genes already known to be involved in human cancer predisposition (*SHROOM2* and *KMT5A* for chromosomes X and 26 respectively) or immunity (*IL17A* for CFA12).

**Figure 6.**
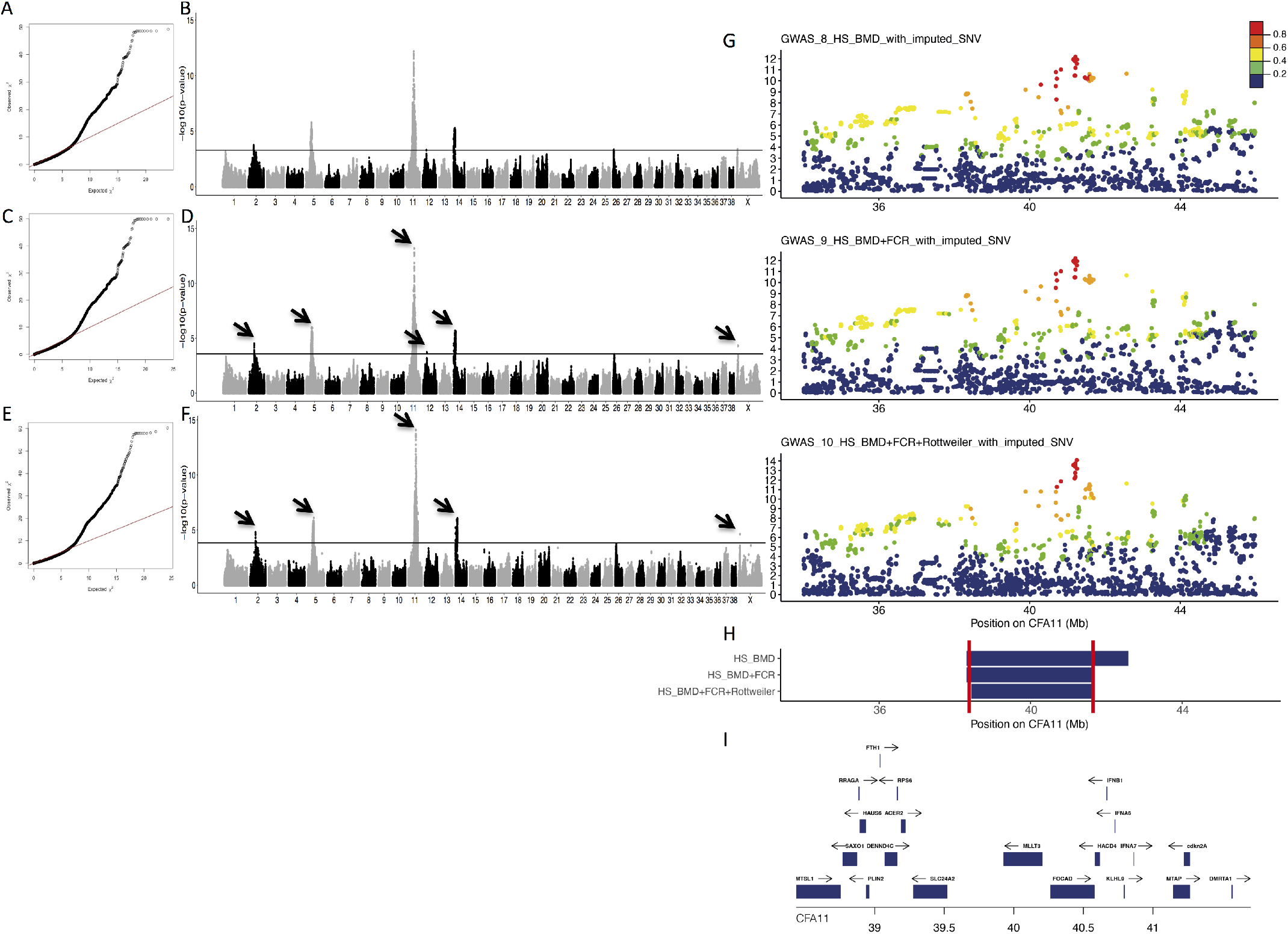
Genome-wide association studies on BMD and other predisposed breeds on HS with the impuation of SNV on a higher density SNV array. A & B: BMD GWAS results based on 404 cases and 371 controls (GWAS_8_HS_BMD_with_imputed_SNV). A) Quantile-Quantile plot displaying a genomic inflation λ of 1.000014, indicating no residual inflation. B) Manhattan plot displaying the statistical results from the GWAS. This analysis pointed out seven loci (arrows) on chromosome 2 (best SNV CFA2: 29716535, *p*_corrected_ = 1.78 x 10^−4^), on chromosome 5 (best SNV Chr5: 30496048, *p*_corrected_ = 1.72 x 10^−6^), on chromosome 11 (best SNV Chr11: 41215628, *p*_corrected_ = 6.65 x 10^−13^), on chromosome 12 (best SNV CFA12: 19862033 *p*_corrected_ = 4.93 x 10^−4^), and on chromosome 14 (CFA14: 8139720, *p*_corrected_ = 5.4 x 10^−6^), on chromosome 26 (best SNV CFA26: 6024801, *p*_corrected_ = 4.7 x 10^−4^) and chromosome X (best SNV CFAX: 6454501, *p*_corrected_ = 4.9 x 10^−4^). C & D: GWAS results for HS combining BMDs (with 404 cases vs 371 controls) and FCRs (13 cases vs 15 controls) (GWAS_9_HS_BMD+FCR_with_imputed_SNV). C) Quantile-Quantile plot displaying a genomic inflation λ of 1.000019, indicating no residual inflation. D) Manhattan plot displaying the statistical results from the GWAS. This analysis pointed out six loci (arrows) on chromosome 2 (best SNV CFA2: 29716535, *p*_corrected_ = 3.09 x 10^−5^), on chromosome 5 (best SNV CFA5: 30496048, *p*_corrected_ = 1.04 x 10^−6^), on chromosome 11 (best SNV CFA11: 41252822, *p*_corrected_ = 6.47 x 10^−14^), on chromosome 12 (best SNV CFA12: 19862033 *p*_corrected_ = 1.99 x 10^−4^), on chromosome 14 (CFA14: 6567456, *p*_corrected_ = 2.06 x 10^−6^) and chromosome X (best SNV CFAX: 6521019, *p*_corrected_ = 4.9 x 10^−4^). E & F: GWAS results for HS combining BMDs (404 cases vs 371 controls), FCRs (13 cases vs 15 controls) and Rottweilers (38 cases vs 22 controls) (GWAS_10_HS_BMD+FCR+Rottweiler_with_imputed_SNV). E) Quantile-Quantile plot displaying a genomic inflation λ of 1.000035, indicating no residual inflation. F) Manhattan plot displaying the statistical results from the GWAS. This analysis pointed out five loci (arrows) on chromosome 2 (best SNV CFA2: 29716535, *p*_corrected_ = 1.59 x10^−5^), on chromosome 5 (best SNV CFA5: 33823740, *p*_corrected_ = 7.89 x 10^−7^), on chromosome 11 (best SNV CFA11: 41252822, *p*_corrected_ = 8.34 x10^−15^), on chromosome 14 (best SNV CFA14: 11021670, *p*_corrected_ = 8.7 x 10^−7^) and on chromosome X (best SNV CFAX: 6521019, *p*_corrected_ = 2.33 x 10^−5^). G) Close up view of the CFA11 locus highlighting the best p-values obtained in the three GWAS experiments and the R^2^ in cases of the SNVs with the top GWAS SNV are depicted to show LD structure: BMDs GWAS (GWAS_8_HS_BMD_with_imputed_SNV), BMDs plus FCRs GWAS (GWAS_9_HS_BMD+FCR_with_imputed_SNV), BMDs plus Rottweilers and FCRs GWAS (GWAS_10_HS_BMD+FCR+Rottweiler_with_imputed_SNV). H) Regions delimitated by SNPs in LD with the best GWAS SNPs (R^2^ >0.6) in cases, the minimal region between the three GWAS (CFA11: 38435917-41701130) is delimitated by red lines. I) Close up view of the genes (with available symbol) located in this 38-42Mb minimal region.

**Table 3:**
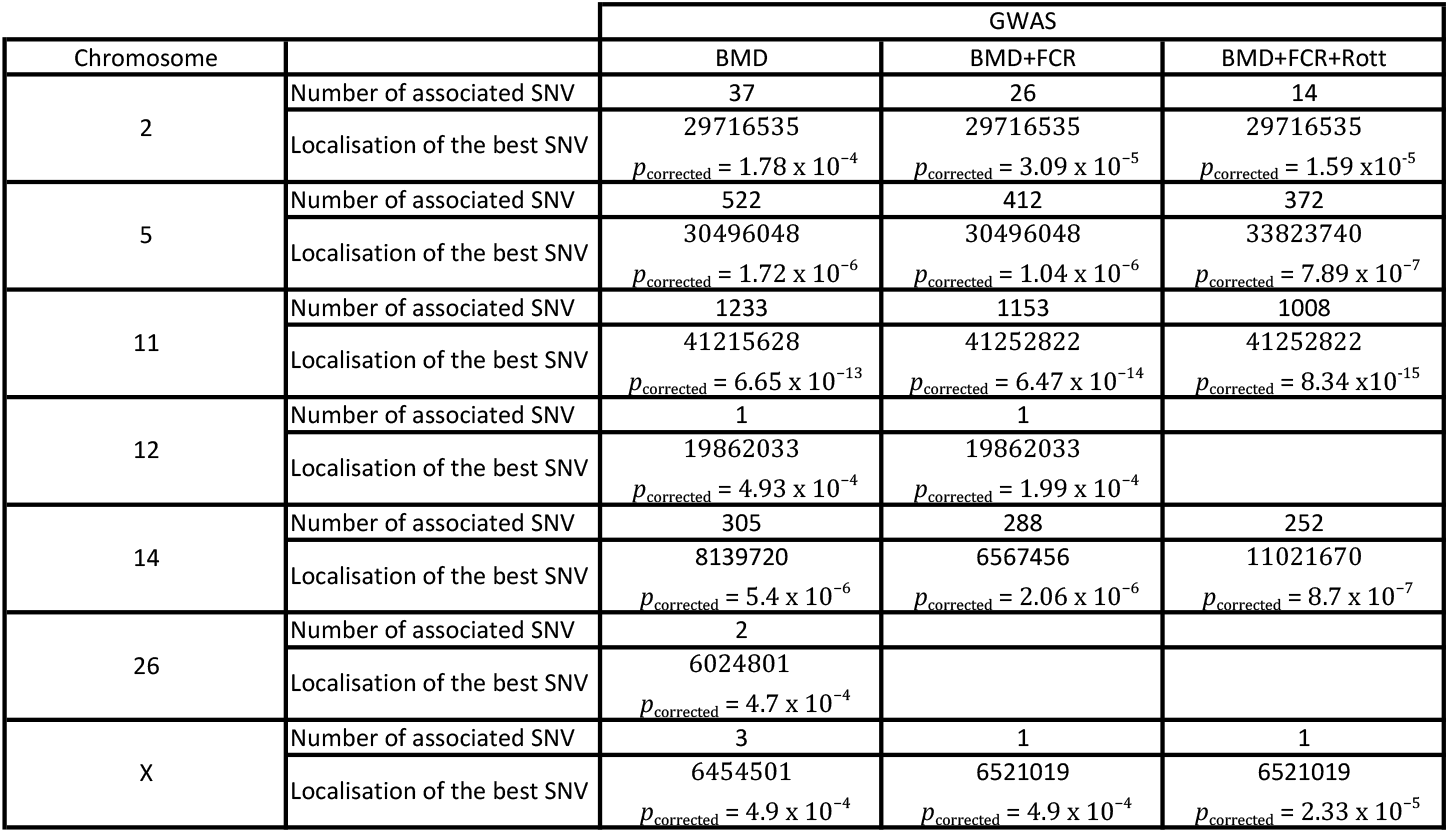
Significant loci identified by GWAS with imputation on high density SNV. Number of linked SNVs with the best SNV and the corresponding corrected p-value are presented for each locus and each GWAS.

The addition of 28 FCRs (13 cases and 15 controls) to previous BMD GWAS for HS with imputed genotyped (GWAS_8_HS_BMD_with_imputed_SNV) identified 1,880 SNVs significantly associated to HS (Table 3) (Figure 6 CD). The increased signal on CFA11: 41Mb with the addition of FCR is in favor of a shared risk locus between FCR and BMD.

The addition of 60 Rottweilers (38 cases and 22 controls) to previous HS BMD and FCR GWAS for HS with imputed genotyped (GWAS_9_HS_BMD+FCR_with_imputed_SNV) identified 1,647 SNVs significantly associated to HS (Table 3) (Figure 6 EF)

These analyses identified HS risk loci shared between the three predisposed breeds on at least chromosomes 2, 5, 11 and 14. A close look at the CFA11 locus showed a large region associated with HS risk (Figure 6 G). Haplotype analysis close to the best CFA11 SNV (41,252822) confirmed that BMD and Rottweiler share the same risk haplotype at position 41 Mb, but no common risk haplotype can been identified with higher SNV density between FCR and other predisposed breeds (Supplementary Table 1). The BMD risk haplotype is frequent in FCR (53.8% and 56,6% in FCR cases and controls) and 75% of FCRs carry at least one copy of this risk haplotype. Yet, while the number of FCR in the GWAS remains low, this haplotype does not appear enriched in HS cases. We identified a second independent CFA11 HS locus between 44 and 45 Mb in GWAS on the three breeds (Figure 6G) and already identified in previous studies^26^. The best SNVs on this region (CFA11:44150645) is not in LD with best SNV of CFA11 (CFA11: 41252822) (R2 on the three breeds: 0.41), confirming that there are at least two independent regions on CFA11 involved in HS predisposition. The haplotype analysis of this CFA11:44Mb region pointed out a common risk haplotype in the three breeds (Supplementary Table 2). Regarding CFA5, the GWAS analysis pointed out a large locus (26-35Mb), overlapping the two CFA5 lymphoma loci previously identified by Tonomura et al (29 Mb and 33Mb)^16^. Similarly, the haplotype analysis around the best SNV (CFA5: 33,823740) pointed out a common risk haplotype delimited by 8 SNVs (34,213158-34,234461) in the 3 breeds and significantly enriched in BMD and FCR cases and common in Rottweilers (69% and 63% in cases and controls, respectively) (Supplementary Table 3). The third main HS risk locus is on CFA14, the haplotype analysis showed that the same risk haplotype is enriched in BMD and Rottweiler cases but not in FCRs cases. Surprisingly, the same haplotype is enriched in controls of the three breeds suggesting that the CFA14 locus contains a protective allele, shared in those predisposed breeds (Supplementary Table 4).

All in all, the imputation with a higher density of SNV allowed the identification of shared risk loci between the 3 HS predisposed breeds (BMD, Rottweiler, FCR). We identified common risk or protective haplotypes shared in predisposed breeds on the major loci localized on CFA11, 14 and 5. In these 3 breeds, cases cumulated risk alleles on these 3 chromosomes, especially in BMD and Rottweilers for which the majority of cases concentrated at least 5 risk copies (72% to 74% of cases vs 31% to 43% of controls, Odds ratio: 5.81, pvalue 1.14×10-31 Chi2 test) (Table Supplementary 5).

### 2.5. Capture and targeted sequencing of the best three HS candidate loci (CFA5, 11 and 14)

Since common haplotypes were detected in the three predisposed breeds on the three main loci (CFA5, CFA11, CFA14), we decided to sequence these three regions to identify putative common variants. The DNAs of 16 dogs (10 BMDs, 4 Rottweilers and 2 FCRs) from the three predisposed breeds with a balanced distribution of risk and protective haplotypes were selected for targeted sequencing. With a mean depth per sample of 142X, 9 458 SNVs and 2 674 Indels per samples were identified. These variants (SNVs and indels) were imputed on the remaining dog samples (455 HS cases and 408 controls from BMD, Rottweiler and FCR breeds). When performing the statistical analysis with the imputed genotypes, no coding variant significantly associated with HS risk could be found and six of the ten top variants associated with HS predisposition are imputed genotypes and are localized within 100 kb on CFA11 (Supplementary Table 6). The best associated variant remains the CFA11: 41,252822 already identified in previous GWAS (§ 3.4). It is localized in a non coding transcript (CFRNASEQ_UC_00018829) which overlaps *CDKN2A* and *CDKN2A-AS*. Interestingly, using RNASeq data from Hoeppner et al^35^, CFRNASEQ_UC_00018829 is highly expressed in the blood tissue compared to 8 others tissues. Moreover the human orthologous region of this SNV (chr9:21 996 622, hg38), closed to CpG island (chr9:21 994 103-21 995 911, hg38) and DNAse I hypersensivity Peak cluster (chr9:21 994 641-21 996 130, hg38), overlaps an enhancer (GH09J021996 chr9:21 996 543-21 996 791, hg38) which regulates *CDKN2A* and *CDKN2B*. The second top SNV is an indel localized in a non coding transcript (CFRNASEQ_IGNC_00021613), 6 500 bp upstream of *CDKN2A* transcript (Supplementary Table 6). Since it was previously shown that the expression of *CDKN2A* correlated with CFA11: 41 Mb risk haplotype^26^, we suspect that these non coding SNVs could act to deregulate the expression of *CDKN2A*.

To identify the best candidate variants in the secondary loci, a complementary association analysis was performed by taking into account the information of CFA11 best SNV (CFA11:41252822) by including the genotypes of this variant as a covariate (Supplementary Table 6). We identified significant residual associations in the three breeds for the CFA11:44-45Mb locus with the best SNV CFA11: 45,941548 (pvalue =0.033), close to C9orf72, not in LD with CFA11: 41,252822 (R2 on the three breeds: 0.0899). This confirmed the existence of two independent risk locus on CFA11 between 44-45Mb.

For the CFA14 locus, six of the ten top variants are imputed variants, localized between position 10951501 and 11043035. This locus overlaps *POT1-AS1* and *POT1*, with the top variant being an indel overlapping *POT1-AS1*.

Regarding the CFA5 locus, eight of the ten top variants of CFA5 found to be associated with the risk to develop HS, are imputed variants; they are localized between position 30488886 and 30489217 overlapping the *SPNS3* gene (Supp Table 6 and Supp Figure 2). Interestingly the second and third variants are localized in a region containing DNA methylation marks, one of them including the CFA5:30489217 variant. Moreover the CFA5:30489203 variant creates a CpG site. We thus hypothesized that these two SNVs could be associated with allele specific methylation in histiocytic cells. Through bisulfite sequencing on histiocytic sarcoma cell lines, we confirmed that these two variants (CFA5:30489203 and CFA5:30489217) presents specific alleles (CFA5:30489203-G and CFA5:30489217-C respectively) creating CpG sites with methylation in histiocytic cells (Supplementary Figure 3). This is also the case for the best HS GWAS SNV on CFA2 with the CFA2:29716535-G allele (Supplementary Figure 4). We hypothesize that these specific methylation alleles could be associated with modifications of the regulation of close genes.

## 3. Discussion

### 3.1. The dog breeds: unique models to detangle the genetic features of human cancers

Through the several genetic studies of canine cancers it has been shown that high risk breeds constitute a major source of advance into the genetics of rare cancers in human^6,8,25^. Indeed, a limited number of critical loci have been identified in canine cancers^6,15–17,26^, with some common loci to several canine cancers and well known in Human cancers too. On the somatic aspects, interestingly, somatic alterations identified to date in canine tumors, even through genome wide approaches, hit the same genes^25,36,37^. Histiocytic Sarcoma, affecting few dog breeds with incredibly high frequencies (BMD, Rottweiler, retrievers) appears as a perfect example to study the genetics underlying such a strong predisposition in dogs, while so rare in humans.

Indeed, dissecting the genetic factors for rare cancers such as histiocytic sarcoma is challenging in Human. We hypothesized that dog models of HS would help to decipher genetic predisposing factors of this rare cancer and indeed, we successfully identified previously a relevant locus on canine chromosome 11, this locus being also well known in canine osteosarcoma and several human cancers^17,38^. Here we present GWAS on a large cohort of several breeds affected by HS. Benefiting from a multiple-breed approach, we not only refined the previously identified locus on CFA11 but also identify additional loci on CFA2, 5, 12, 14, 26, 20 and X. Moreover, this study highlights the fact that behind an initial association signal peak found in canine cancer GWAS, independent risk haplotypes can be cumulated and shared with several dog breeds and several cancers.

### 3.2. The genetic predisposition of canine cancers: cumulative of risk haplotypes

In Humans, the genetic architecture of cancer risk is usually described as a combination of rare variations in families with dominant inheritance patterns, and common variants with small effect sizes in the population at large. While GWAS in dogs, in a given breed, usually described fewer variants with stronger effects^18^ due to breed structure and artificial selection of variants with strong effects, which is not necessarily the rule for cancer. Indeed, due to the genetic drift of canine breeds due to the strong selection of sires and dams used for reproduction on beauty selection criteria, deleterious alleles could be involuntary selected and enriched in specific populations. As a consequence, significant associations detected with cancer in dog GWAS could be due to cumulative risk alleles. In those conditions, it would be surprising that HS predisposition was only due to one risk haplotype, especially since breeders did not manage to reduce prevalence and with at least 33 loci that have been identified by GWAS in canine osteosarcoma predisposition^17^. Our results confirmed the main role of CFA11 locus with at least the cumulative effect of two different risk haplotypes. The existence of 2 close loci was already suggested in our previous publication^26^. The present work, using a multi-breed approach. Moreover this work allowed the identification of a strong candidate variant overlapping *CDKN2A-AS* regulating *CDKN2A*. Additional GWAS peaks were identified on CFA2, 5, 11, 12, 14, 20, 26 and X. Some of these loci are shared between the 3 predisposed breeds, mostly between Rottweiler and BMD which is expected considering close genetic relationship between these two breeds^39^. Considering the 3^rd^ predisposed breed, due to the fact that the FCR display small numbers of dogs in France, thus the number of FCR samples included in the study is low and further GWAS will be needed to better decipher shared predispositions between FCR and other breeds. Nevertheless, in these three predisposed breeds (BMD, Rottweiler and FCR), the cumulative risk alleles on the 3 main loci (CFA11, 5, 14) strongly impact the probability to develop this cancer with an Odds ratio of 5.41 to be affected by HS when dogs carry 5/6 risk alleles. This work illustrates that the GWAS association detected between a cancer and a locus in dogs could hide the cumulative risk of several haplotypes. This point is confirmed by Arendt et al. or Tonomura et al who described in the golden retriever breed at least 2 independent risk haplotypes on the CFA20 locus for mast cell tumor and on the CFA5 locus for lymphoma respectively^15,16^.

### 3.3. Common risk haplotypes to several cancers identified through a multi-breed approach

On the other hand, the present study confirms that some risk haplotypes are also involved in several cancers, as suggested by Tonomura et al. with the association of the CFA5 locus with hemangiosarcoma and lymphoma^16^. Such pleiotropy at cancer risk loci has also been observed in human cancers, for which one-third of SNVs map to genomic loci associated with multiple cancers^38^. Here we confirmed the pleiotropic effect of loci for CFA5, CFA11 and CFA20 loci, influencing the risk for HS and lymphoma, osteosarcoma or mast cell tumor in BMD or golden retriever. Further studies, with higher SNV density in the golden retriever breed are needed to confirm whether or not the same predisposing risk alleles are shared with BMD. Indeed in human GWAS, for some loci, different risk SNVs are associated to different risk cancers, though they might ultimately converge to the same oncogenic mechanism^38^. Regarding the *CDKN2A* locus, detected in Rottweiler and BMD, interestingly a neighboring region of the locus (CFA11: 41.37MB CanFam3) was found to be associated to osteosarcoma and is fixed in the Rottweiler population^17^. The osteosarcoma risk haplotype is fixed in our Rottweiler population showing that HS affected Rottweilers are cumulating risk haplotypes for at least 2 cancers (Osteosarcoma and HS) at this locus. This co-occurrence of 2 different risk haplotypes for HS and osteosarcoma across 200kb also perfectly illustrate that a given locus can harbor two risk haplotypes for two different cancers. In this condition, using GWAS in a multi-breed strategy should help to decipher risk haplotypes when several breeds share several predispositions.

### 3.4. Pleiotropic effect of loci

Cancers are multigenic diseases with cumulative alterations in key pathways considered as hallmarks of cancer^40^. HS involving histiocytic immune cells is suspected to be at the crossroads of immune dysregulation and cancer predispositions in dogs. Indeed HS predisposed breeds, especially BMD, are also predisposed to reactive histiocytic diseases^41^ but also to other immune or inflammatory diseases such as glomerulonephritis, aseptic meningitis or inflammatory bowel disease (https://www.bmdca.org/health/diseases.php)^42,43^. While no causal relationship was proved between inflammation and HS, inflammation is suspected to contribute to HS development^44–46^. In these conditions it is not surprising that HS GWAS hits overlap not only candidate tumor suppressor genes (*TUSC1*) or well-known tumor suppressors involved in cell cycle (*CDKN2A*), in genome stability (telomere protection: *POT1*, replication stress/DNA damage: FHIT…) but also in inflammation (*IL17a, IL17rd, SPNS3, ARHGEF3*…) (Supplementary Table 7). The GWAS hits highlight an enrichment of genes involved in the IL17 pathway (pvalue: 0.0014204), pathways of cancers (G1 to S cell cycle control -pvalue 0.0055919-, Non-small cell lung cancer -pvalue 0.0070335-, Signaling Pathways in Glioblastoma -pvalue: 0.0090479-, Small cell lung cancer -pvalue 0.012008, TP53 Network -pvalue 0.033586-) and lipid metabolism (Oxysterols derived from cholesterol -pvalue 0.031844-, Leptin and adiponectin -pvalue 0.017808-). Moreover, we expanded our search of overlap between the HS association signals in this study and human GWAS signals (Supplementary Table 7). A number of these genes are not only already known to be involved in the predisposition of several cancers (*CDKN2A*, *POT1*, *FHIT*,.) but they are also associated with immune traits (monocyte, platelet.) or with cholesterol, HDL/LDL or allergens traits in Human. This result suggests that the pleiotropic nature of these loci is not limited to cancer risk, as shown in human, with the fact that loci associated with N-glycosylation of human immunoglobulin G show pleiotropy with autoimmune diseases and haematological cancers^47^. Concomitantly to this work, Labadie et al study confirmed the pleiotropy effect of these cancers loci identified in canine breeds, with shared region for canine T zone lymphoma, mast cell tumour and hypothyroidism in golden retriever; one of these loci on CFA 14 involved in mast cell tumour and canine T zone lymphoma is ~800kb downstream of *POT1* locus identified in this study^48^.

### 3.5. The genetic predisposition of Histiocytic sarcoma in 3 dog breeds (CFA 5, 14 and 20)

In addition to the CFA11 major locus, this study clearly identified other HS loci on CFA5, 14 and 20. The locus of CFA5 is associated with HS and lymphoma risk in BMD. The top CFA5 locus SNVs are located in an intron of the *SPNS3* gene (sphingolipid transporter 3). Very few is known about this gene although its paralogous gene (*SPNS2*) is known to be important in immunological development, playing a critical role in inflammatory and autoimmune diseases, influencing lymphocyte trafficking and lymphatic vessel network organization and driving defective macrophage phagocytic function (https://www.ncbi.nlm.nih.gov/gene/124976,^49^). Moreover this region is associated in human GWAS with the chemokine CLL2. Thus, *SPNS3* is a strong candidate gene to explain HS and lymphoma predispositions.

For the locus on CFA 14, the best SNVs are located in introns of *POT1-AS1* and *POT1. POT1* gene encodes a nuclear protein involved in telomere maintenance and is involved in the predisposition and the development of numerous cancers (supp table 7). This locus was suggested in our previous work^26^. This study identified a potential protective haplotype shared between the three predisposed breeds. The difference of the age of onset of cases carrying zero or one copy, as well the difference of the age of death of controls carrying none/one or two copy of this protective haplotype suggests that this shared protective haplotype is most probably involved in longevity and thus also in the age of onset of HS (Supplementary Figure 5).

The locus of CFA20 is associated with HS and mast cell tumor risk in BMD. The top SNV of CFA20 locus is intronic of *FHIT* a tumor suppressor involved in apoptosis and prevention of the epithelial-mesenchymal transition, one of earliest and most frequently altered genes in the majority of human cancers^34^, and involved in human predisposition of breast cancer. Moreover the stable nuclear localization of FHIT is a special marker for histiocytes suggesting an other function for FHIT as a signaling molecule related to anti-proliferation function^50^.

### 3.6. Causal variants

As expected with previous work on canine HS predisposition^26^ and human cancers^38,51^ capture and targeted sequencing of the best three HS candidate loci did not allow to identify potential coding variants in these three loci and the risk variants are common in the predisposed breeds. Like in human, the majority of the loci identified from cancer GWAS do not directly impact on the amino acid sequence of the expressed protein and the elucidation of causal variants is challenging, since all closely linked variants that are in LD with the best GWAS SNV are relevant candidates^38^. In this study, best SNVs are either in intronic part of candidate genes (*FHIT*, *SPNS3*,.), upstream of the candidate gene (*POT1*) or overlapping non coding RNA in the vicinity of strong candidate gene (*CDKN2A, POT1*). Only one variant associated to HS risk in FCR and BMD was found in the 3’ UTR of *IL17a*. Finally, we showed that the best variants linked to *SPNS3* or *PFKFB3*, belong to CpG sites methylated in histiocytic cells. All these elements strongly suggest that predisposing variants to HS in dogs are non coding variants, with regulatory effects. In those conditions, the local cumulative risk haplotypes could reflect complex regulatory interactions like for the *MYC* locus, for which recent Hi-C analysis of its locus has demonstrated a more complicated regulatory mechanism, implicating various large intergenic non-coding RNAs that mediate effects at risk loci^38^. Further functional studies are needed to identify the involvement of such variants on the regulation of candidate genes.

To conclude, we present here the largest GWAS of histiocytic sarcoma in dog cohorts, through a multi-breed approach, confirming the main role of *CDKN2A* locus (CHA11/HSA9) and identifying three new loci (CFA5, 14 and 20). With multiple breeds and cancers approach, we highlight the cumulative effect of different risk haplotypes behind each locus and their pleiotropic nature.

## 4. Materials and Methods

### 4.1. Sample Collection

Blood and tissue biopsy samples from cancer affected and unaffected dogs were collected by a network of veterinarians through the Cani-DNA BRC (http://dog-genetics.genouest.org) and DNA/RNA were extracted as previously described [Ulve et al.]. The work with dog samples was approved by the CNRS ethical board France (35-238-13). The samples were collected by veterinaries in the course of the medical cares of the dogs, with their owner consent: blood and tissue were collected at medical visit or at surgery, then stored in tubes containing EDTA or RNAlater, respectively.

### 4.2. Genome-Wide Association Study (GWAS)

DNAs were genotyped on the Illumina 173k SNV Canine HD array at CNG (Evry, France) and on the Affymetrix Axiom Canine Genotyping Array Set A and B 1.1M SNV array at Affymetrix, Inc. (Santa Clara, USA). We used mixed linear model analyses, taking into account population structure and kinship (Eigenstrat – GenABEL 1.8)^52^, removing SNV with a minor allele frequency under 0.01. P-value corrected for inflation factor λ were used and the threshold for genome-wide significance for each association analysis was defined based on 500 permutations.

For imputation, the Beagle software^53^ [version 4.1] was used to impute the Illumina 20k SNV and Illumina 170k SNV arrays to the Affymetrix 712k SNV array format, as well as to input 21,614 variants identified from the capture and targeted sequencing of 16 dogs. All default settings of the Beagle software were used except for the following options: *niterations=50 window=3000*.

### 4.3. Capture and sequencing of targeted canine GWAS loci

In total, four loci were captured and sequenced for 16 dogs selected according to their haplotype. One localized on CFA5 (29 805 467-34 459 320), two on CFA11 (41 148 019-41 237 204 and 43 520 875-46 778 525) and one on CFA14 (9 949 911-11 524 424).

Capture, sequencing, variants detection and annotation were done by IntegraGen S.A. (Evry, France). Genomic DNA was captured using Agilent in-solution enrichment methodology, i.e. Agilent SureSelect Target Enrichment System kit (Agilent technology, Santa Clara, California, USA). The SureSelect Target Enrichment workflow is a solution-based system using ultra-long – 120-mer – biotinylated cRNA baits – to capture regions of interest, enriching them out of a NGS genomic fragment library. The library preparation and capture has been followed by paired-end 75 bases massively parallel sequencing on a Illumina HiSeq 2000 sequencer. For detailed explanations of the process, see Gnirke et al.^54^.

A custom-made SureSelect oligonucleotide probe library was designed to capture the loci of interest according to Agilent’s recommendations with a 1X and 2X tiling density, using the eArray web-based probe design tool (https://earray.chem.agilent.com/earray). A total of 57,205 RNA probes were synthesized by Agilent Technologies, Santa Clara, CA, USA.

Sequence capture, enrichment and elution were performed according to manufacturer’s instruction and protocols (SureSelect, Agilent) without modification except for library preparation performed with NEBNext^®^ Ultra kit (New England Biolabs^®^). For library preparation 600 ng of each genomic DNA were fragmented by sonication and purified to yield fragments of 150-200 bp. Paired-end adaptor oligonucleotides from the NEB kit were ligated on repaired A tailed fragments, then purified and enriched by 8 PCR cycles. 1200 ng of these purified libraries were then hybridized to the SureSelect oligo probe capture library for 72 hr. After hybridization, washing, and elution, the eluted fraction was PCR-amplified with 9 cycles, purified and quantified by QPCR. Based on this quantification, an equimolar pool was performed then quantified again by QPCR. Finally, the pool was sequenced on an Illumina HiSeq 2000 as paired-end 75b reads. Image analysis and determination of the bases were made using Illumina RTA software version 1.12.4.2 with default settings.

The bioinformatic analyses of sequencing data were based on the Illumina pipeline (CASAVA1.8.2). CASAVA performs alignment of a sequencing run to a reference genome (canFam3), calls the SNVs based on the allele calls and read depth, and detects variants (SNVs and Indels). The alignment algorithm used was ELANDv2 (Maloney alignment and multi-seed reducing artifact mismatches). Only the positions included in the bait coordinates were conserved. Genetic variation annotation was performed by IntegraGen in-house pipelines. It consists on genes annotation (RefSeq), detection of known polymorphisms (dbSNP) followed by variant annotation (exonic, intronic, silent, nonsense..).

### 4.4. Haplotype analyses

Minimal risk haplotypes for different breeds were identified on the CFA11 associated locus. Firstly, variants in strong LD (r2>0.8) with the top SNV were identified in each breed using PLINK 1.9^55^ LD clumping and used as input for haplotype phasing in each breed with fastPHASE^56^ 1.4. The risk haplotypes enriched in cases were identified based on the top SNV genotype. Starting from top SNV localization and walking both up- and downstream, we then identified the SNV-positions where the risk haplotype was broken by a recombination event (i.e. two alternative alleles were present on both risk and non-risk haplotypes). This was done separately for each breed, and thereafter the minimal shared risk haplotype across breeds was defined.

### 4.5. Gene-set enrichment analysis

Approved symbols of the closest genes to the strongest significant signal per chromosome and per GWAS were analysed with online tool Webegestalt^57^ to identify over-representation of pathway in the loci of the GWAS.

### 4.6. Methylation analyses

Methylation analyses were performed on 12 histiocytic sarcoma cell lines, including one commercial cell line (DH82; ATTC, CRL-10389; RRID:CVCL_2018) and eleven cell lines developed in the team from HS affected dog fresh tissues. These eleven cell lines are available on request. Cells were cultivated in a complete RPMI medium containing RMPI 1640 GlutaMAX supplemented medium (Gibco. life technologies) with 10% of Fetal Bovine Serum (HyClone, GE Healthcare, Life Sciences, Logan, UT) and 0.025% of Primocin (InvivoGen, Toulouse France) at 37 C in a humidified 5% CO2 incubator. All cell lines were tested for mycoplasma with MycoAlertTM Plus kit (Lonza, Rockland, ME) and were mycoplasma-free cells. The SNVs of CFA5 and CFA2 were sequenced by Sanger sequencing as previously described^10^ with the following primers: CFA2_29716535-F: GGTGTACTTTCGGGTCCAAC, CFA2_29716535-R:CCCTGTCATTCGATGTCCTT, CFA5_30489203-30489217_F: CCTGAGTGAGTGGAATGAGGA, CFA5_30489203-30489217_R: CTTCCTGCGACCTGCTGT) in absence and or (CFA2_29716242-29716795_FM: TAGGTGTTGGGTTTATATTGTTAGG, CFA2_29716242-29716795_RM: CTTCCTGCGACCTGCTGT, CFA5_30488738-30489339_FM: TAGGTGTTGGGTTTATATTGTTAGG CFA5_30488738-30489339_RM) in presence of bisulfite conversion (EZ DNA Methylation-Gold Kit, Ozyme, St Cyr -l’école, France).

## Acknowledgments

We thank veterinarians who provided anatomopathologic diagnoses, especially Olivier Albaric and Laetitia Dorso (Laboniris, Oniris, Ecole Nationale Vétérinaire de Nantes, France) as well as Marie-Odile SEMIN (LAPV, Amboise, France), Caroline Laprie (Vet-Histo, Marseille, France), Marie Lagadic (Idexx Alfort, France) and Frédérique Degorce-Rubiale (LAPVSO, Toulouse, France). We thank the veterinarians for providing us with clinical data and samples, as well as dog owners, breeders and breed clubs, especially the French club AFBS, European clubs, the American club BMDCA, the IWG International Working Group (for Bernese Mountain Dogs) and the US Berner garde foundation, especially Pat Long for her dedicated trust and follow-up of our work. We also warmly thank Clotilde de Brito, Laetitia Lagoutte, Annabelle Garand, Anne Sophie Guillory, Melanie Rault and Ronan Ulve (IGDR, Rennes, France) for their help to sample collection and Stéphane Dréano (IGDR, Rennes, France) for Sanger sequencing and the Biogenouest bioinformatic platform. The authors thank G. Queney, A. Thomas and C. Dufaure de Citre from Antagene (Animal Genetics company, Lyon, France) for providing samples and genetic sample characterization.

## Supports

This study was supported by CNRS and in the frame of the French Plan Cancer 2009-2013, by INCa PLBio “canine rare tumours” funding (N° 2012-103; 2012-2016) and by Aviesan/INSERM MTS 2012-06, as well as by the American Kennel Club Canine Health foundation, AKC CHF funding N°2446. The sample collection performed through the French Cani-DNA BRC was funded by the CRB-Anim infrastructure ANR-11-INBS-0003 in the frame of the ‘Investing for the Future Program (PIA1). Last, donations from Bernese Mountain dogs Associations were kindly offered by the French AFBS, the Italian SIBB and the German DCBS.

---

**Supplementary Figure 1:** Genome-wide association studies (GWAS) on HS and lymphoma. A & B: BMD GWAS results for HS with 172 cases and 154 controls (GWAS_1_HS_BMD). A) Quantile-Quantile plot displaying a genomic inflation λ of 1.000005, indicating no residual inflation. B) Manhattan plot displaying the statistical results from the GWAS. This analysis pointed out three loci (arrows) on chromosome 5 (CFA5: 30496048, *p*_corrected_ = 6.36 x 10^−5^), on chromosome 11 (CFA11: 41161441, *p*_corrected_ = 4.85 x 10^−7^) and on chromosome 20 (CFA20: 30922308, *p*_corrected_ = 2.52 x 10^−5^). C & D: BMD GWAS results for HS and lymphoma with 252 cases vs 146 controls (GWAS_2_HS+lymphoma_BMD). C) Quantile-Quantile plot displaying a genomic inflation λ of 1, indicating no residual inflation. D) Manhattan plot displaying the statistical results from the GWAS. This analysis pointed out two loci (arrows) on chromosomes 11 (CFA11: 41161441, *p*_corrected_ = 3.18 x 10^−6^) and 5 (CFA5: 30496048, *p*_corrected_ = 3.13 x 10^−6^). E & F: Meta-analysis combining the BMD GWAS for HS and lymphoma (252 cases vs 146 controls) and the golden retrievers GWAS for lymphoma (41 cases vs 172 controls) from Tonomura *et al*. 2015 (GWAS_3_HS+lymphoma_BMD+golden_retriever). E) Quantile-Quantile plot displaying a genomic inflation λ of 1.000008, indicating no residual inflation. F) Manhattan plot displaying the statistical results from the GWAS. This analysis pointed out the locus on chromosome 5 (CFA5: 33001550, *p*_corrected_ = 8.96×10^−8^).

**Supplementary Figure 2.** Genome-wide association studies (GWAS) on HS and mast cell tumor. A & B: BMD GWAS results for HS with 172 cases and 154 controls (GWAS_1_HS_BMD). A) Quantile-Quantile plot displaying a genomic inflation λ of 1.000005, indicating no residual inflation. B) Manhattan plot displaying the statistical results from the GWAS. This analysis pointed out three loci (arrows) on chromosome 5 (CFA5:30496048, *p*_corrected_ = 6.36 x 10^−5^), on chromosome 11 (CFA11:41161441, *p*_corrected_ = 4.85 x 10^−7^) and on chromosome 20 (CFA20:30922308, *p*_corrected_ = 2.52 x 10^−5^). C & D: BMD GWAS results for HS and mast cell tumor with 216 cases vs 135 controls (GWAS_4_HS+MCT_BMD). C) Quantile-Quantile plot displaying a genomic inflation λ of 1.000014, indicating no residual inflation. D) Manhattan plot displaying the statistical results from the GWAS. This analysis pointed out three loci (arrows) on chromosomes 11 (CFA11:41161441, *p*_corrected_ = 1.31 x 10^−6^), 18 (CFA18: 45622322, *p*_corrected_ = 1.52 x 10^−5^) and 20 (CFA20: 30922308, *p*_corrected_ = 4.05 x 10^−6^). E & F: Meta-analysis combining the BMD GWAS for HS and mast cell tumor (216 cases vs 135 controls) and European golden retrievers GWAS for mast cell tumor (65 cases vs 62 controls) from Arendt et *al*. 2015 (GWAS_5_HS+MCT_BMD+golden_retriever). E) Quantile-Quantile plot displaying a genomic inflation λ of 1.000013, indicating no residual inflation. F) Manhattan plot displaying the statistical results from the GWAS. This analysis pointed out the locus on chromosome 20 (CFA20:33321282, *p*_corrected_ = 2.36×10^−7^).

**Supplementary Figure 3**. Identification of DNA methylation sites at the SNVs on the CFA5 locus included in a CpG islands. The UCSC track of chr5: 30,489,183-30,489,230 with Methylation track (Dog-MDCK-Meth) and the Sanger sequencing performed on histiocytic sarcoma cell lines were represented. The sequencing of homozygous and heterozygous histiocytic sarcoma cell lines in absence and presence of bisulfite treatment showed that the two SNVs present allele-specific methylation in histiocytic cells.

**Supplementary Figure 4**. Identification of DNA methylation sites at the SNVs on the CFA2 locus included in a CpG islands. The UCSC track of chr2: 29,716,519-29,716,550 with Methylation tracks (Dog-MDCK-Meth, Dog-R3-Sperm-Meth) and the Sanger sequencing performed on histiocytic sarcoma cell lines were represented. The sequencingof homozygous and heterozygous histiocytic sarcoma cell lines in absence and presence of bisulfite treatment showed that the SNV presents allele-specific methylation in histiocytic cell lines.

**Supplementary Figure 5**. Age of onset or death of BMD HS cases and controls according to the number of copy of the protective CFA14 haplotype. Cases with zero copy - n: 183 - Mean age: 6.22. Cases with one copy - n: 24 - Mean age: 7.23. Cases with one copy - n: 1 - Mean age: 6.9. Controls with zero copy - n: 132 - Mean age: 11.21. Controls with one copy - n: 50 - Mean age: 11.27. Controls with one copy - n: 5 - Mean age: 11.97. Wilcoxon rank sum test, one.sided).

