## Supplementary figures and images for "The tree that hides the forest: identification of common predisposing loci in several hematopoietic cancers and several dog breeds"

### Supplementary figure 1

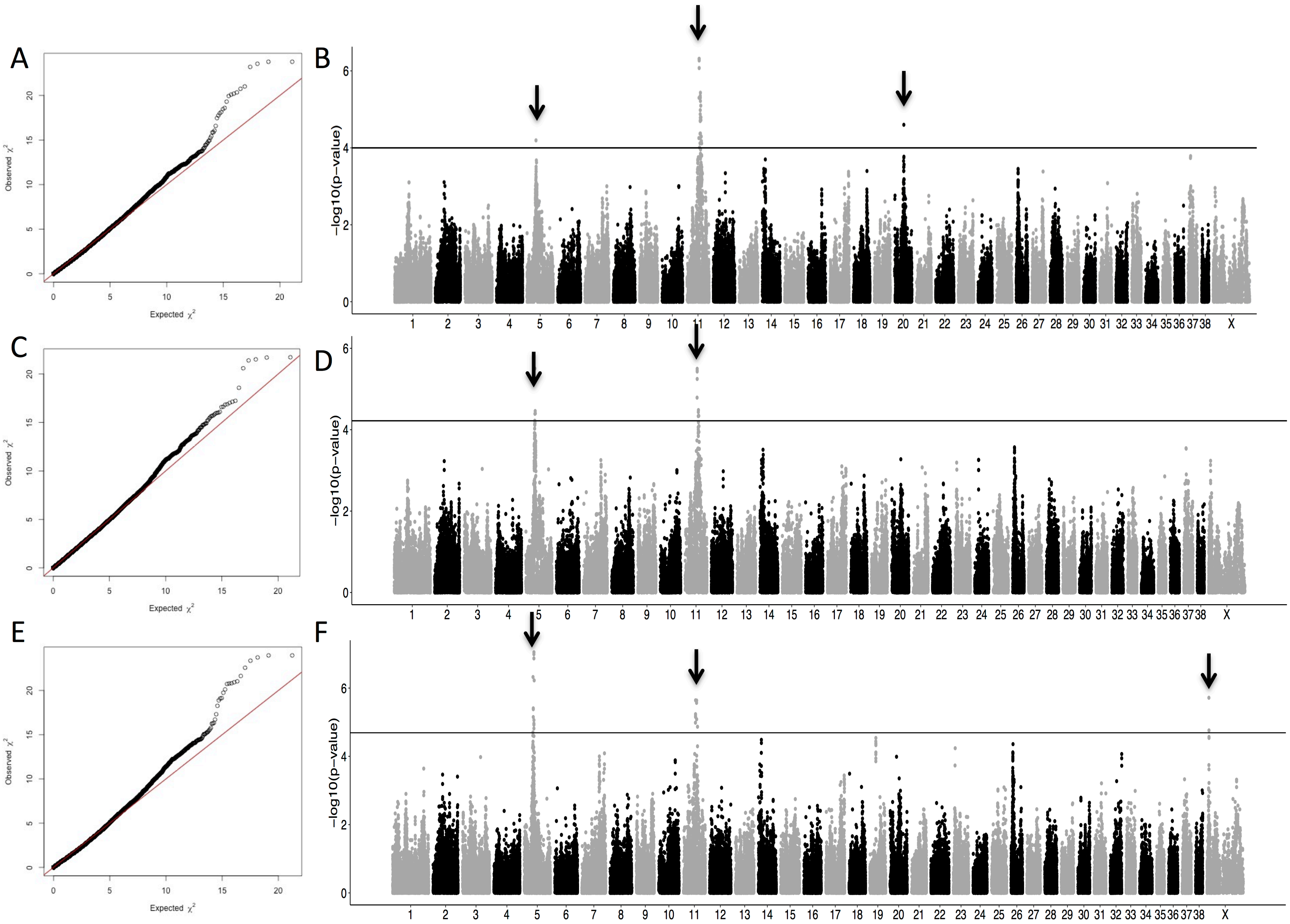

### Supplementary figure 2

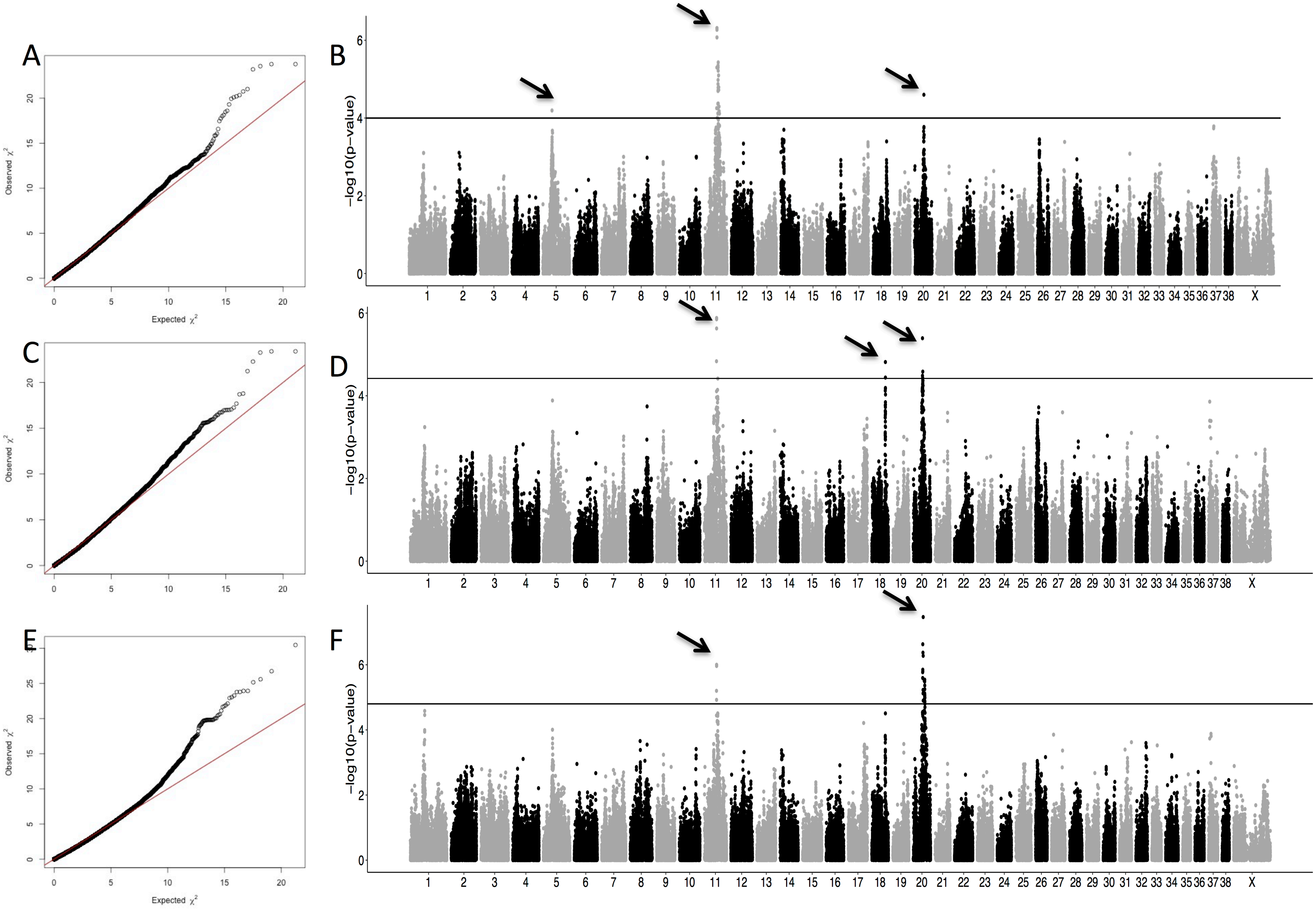

### Supplementary figure 3

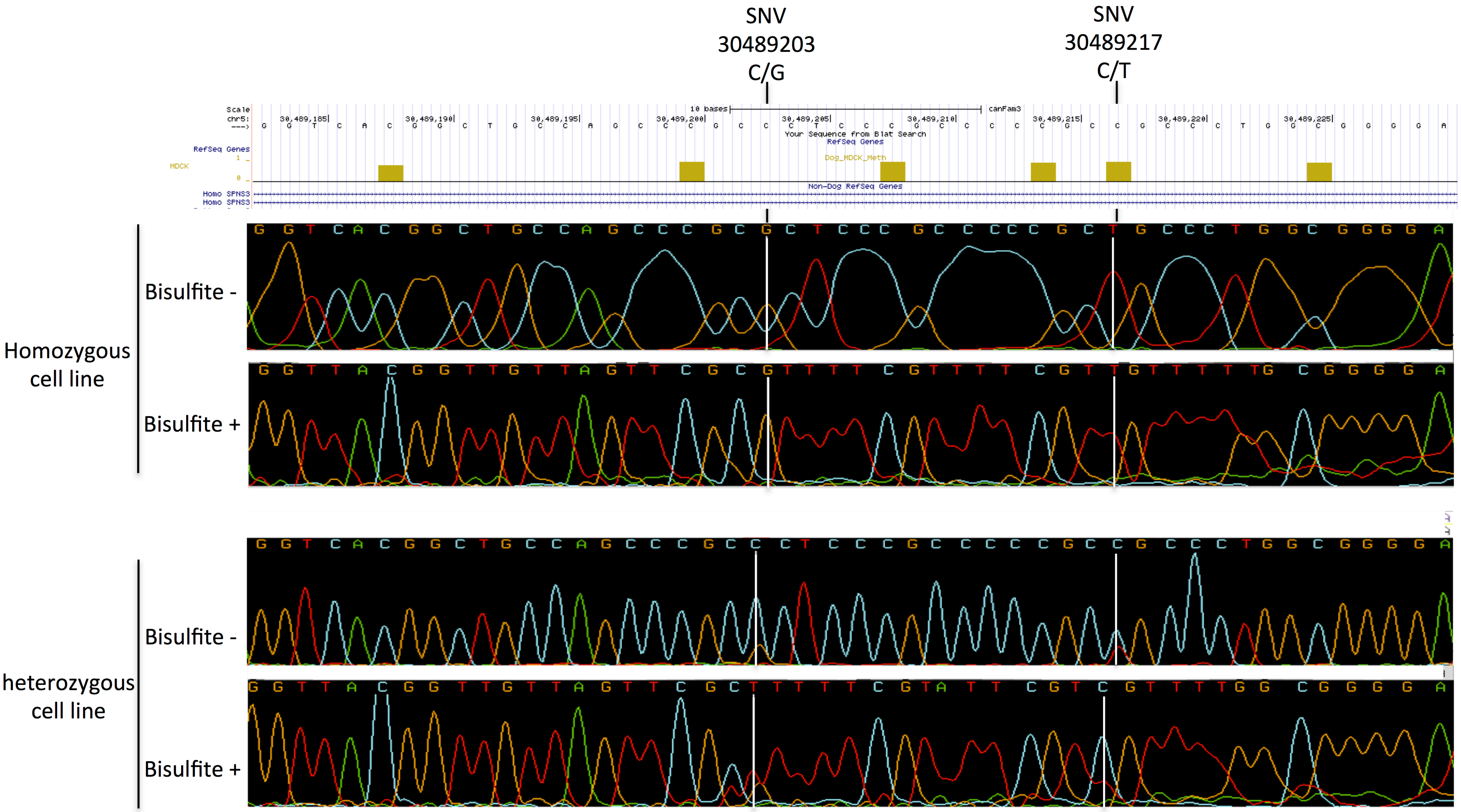

### Supplementary figure 4

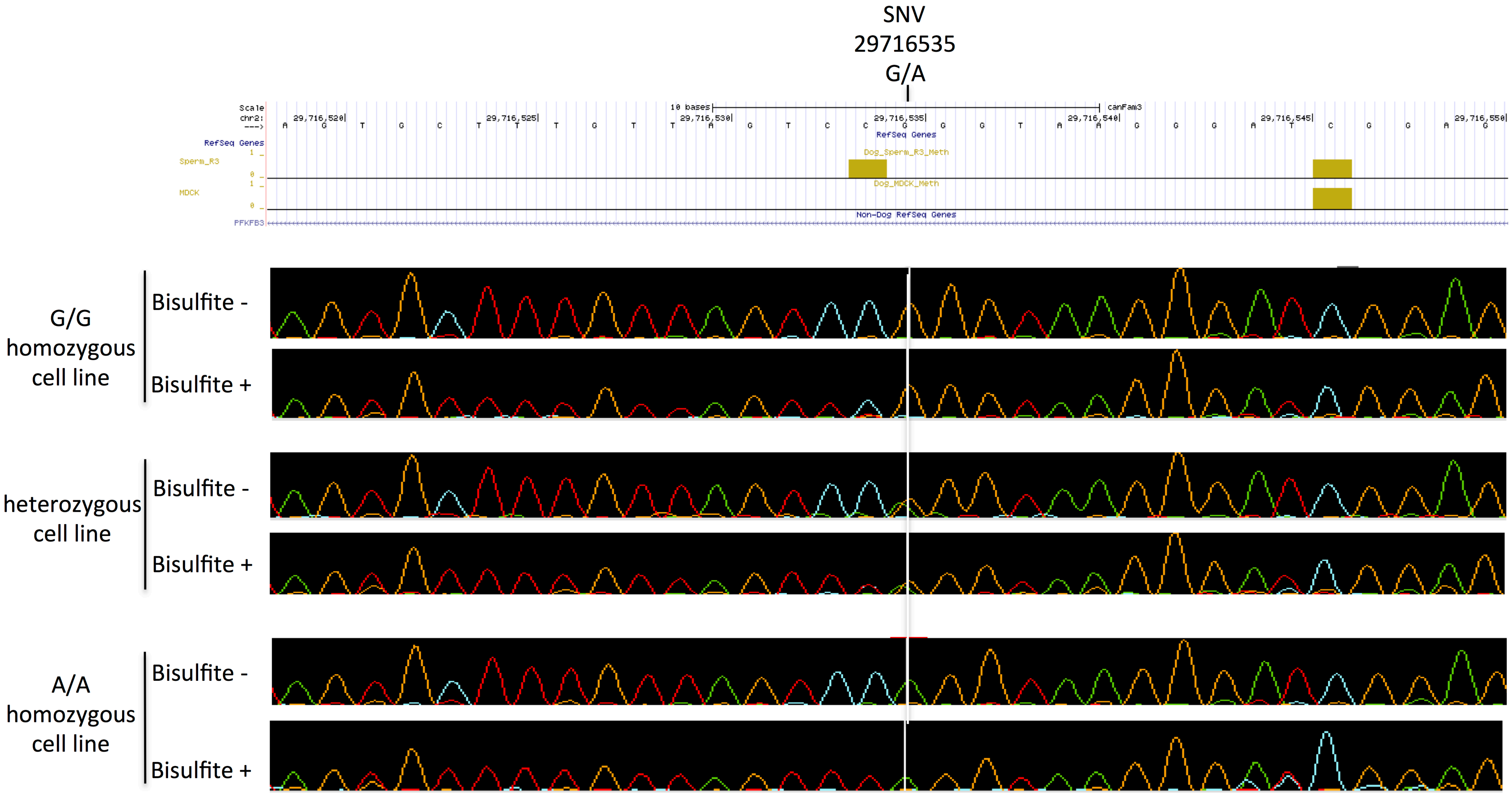

### Supplementary figure 5

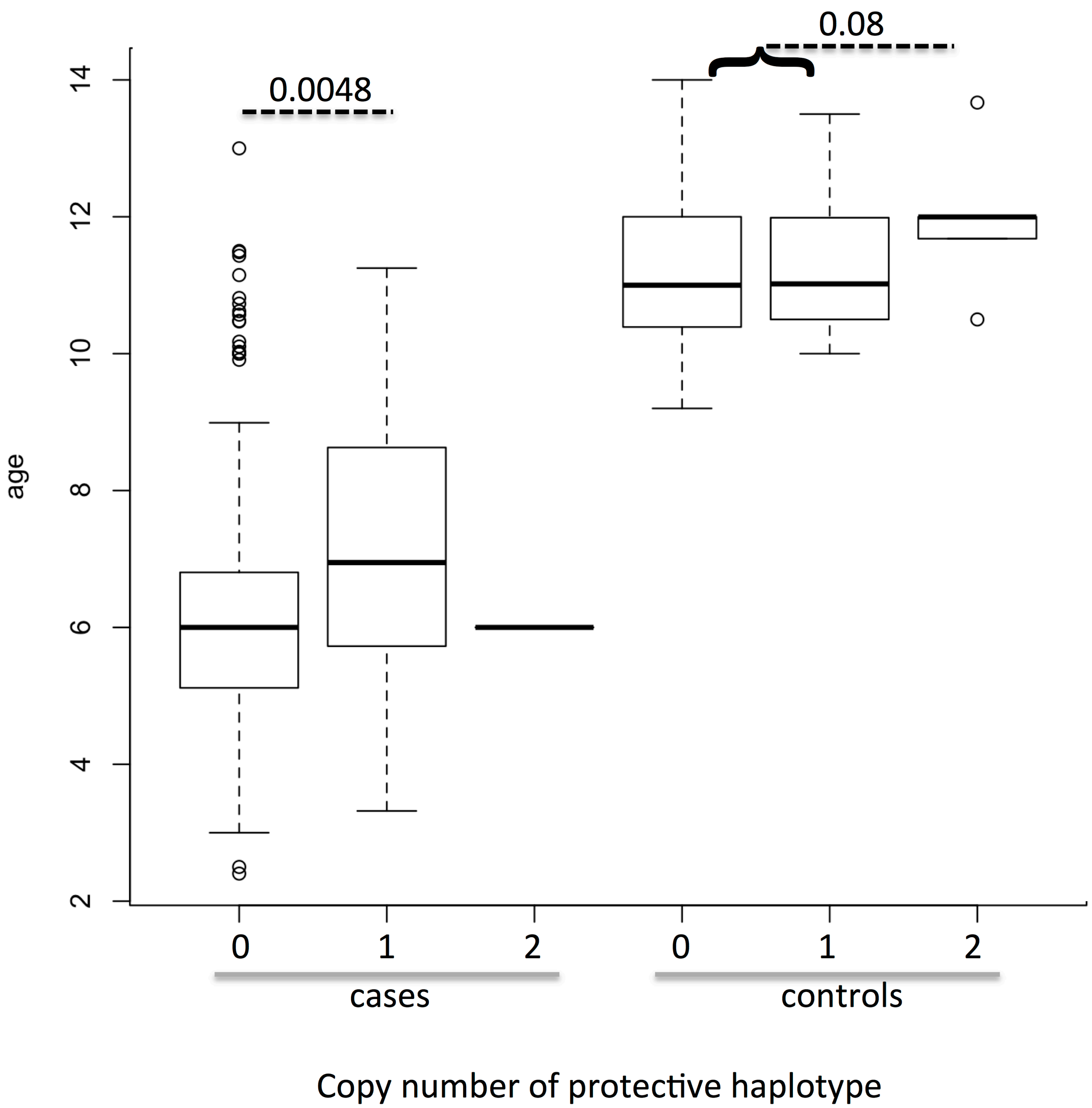
