## Supplementary Table 2 for "The tree that hides the forest: identification of common predisposing loci in several hematopoietic cancers and several dog breeds"

Supplementary table 2: Association analysis of haplotype of CFA11: 44Mb region with the phenotype in predisposed breeds.
The haplotype was determined by the genotype of the following SNVs (43263273, 43272660, 43290639, 43317981, 43690075, 44070751, 44097553, 44098601, 44099662, 44121253, 44126921, 44129025, 44135605, 44150645, 44152497, 44364615, 44366678, 44367106). At risk haplotype in the breeds are represented in bold. CI: confidence interval (Woolf Method).

| | Haplotype frequencies in 3 breeds | | | Odds Ratio | | |
|---|---|---|---|---|---|---|
| | haplotype | affected | unaffected | Odds Ratio | confidence interval (Woolf Method) | pval |
| BMD | **ATGATAGGAACGGCAACT** | 0,65 | 0,44 | 2.39 | [1.95-2.93] | 3.82X10-17 |
| | TCAGTTATTGTAATCGTC | 0,12 | 0,25 | 0.42 | [0.32-0.55] | 9.74X10-11 |
| | TCAGTTATTGCGGTCGTC | 0,09 | 0,13 | 0.66 | [0.48-0.92] | 0.013 |
| | TCAGCAATTGCGGTCGTC | 0,09 | 0,11 | 0.82 | [0.59-1.14] | 0.24 |
| | others | 0,05 | 0,08 | 0.59 | [0.39-0.9] | 0.015 |
| Rottweiler | **ATGATAGGAACGGCAG{T/C}C** | 0,82 | 0,57 | 3.52 | [1.55-8] | 0.002 |
| | ATGACTATTGTAATCGTC | 0,08 | 0,17 | 0.4 | [0.13-1.24] | 0.099 |
| | ATGATAATAGCGGCAACT | 0,05 | 0,15 | 0.3 | [0.08-1.09] | 0.097 |
| | others | 0,05 | 0,11 | 0.44 | [0.11-1.73] | 0.28 |
| FCR | **ATGATAGGAACGGCAA{T/C}T** | 0,35 | 0,17 | 2.65 | [0.76-9.29] | 0.137 |
| | TCAGCAGGAACGGCAATC | 0,54 | 0,7 | 0.5 | [0.17-1.5] | 0.21 |
| | others | 0,12 | 0,13 | 0.85 | [0.17-4.2] | 1 |
| | | 1 | 1 | | | |
| all breeds | **ATGATAGGAACGGCA---** | 0,66 | 0,44 | 2.45 | [2.02-2.98] | 9.47x10-20 |
| | others | 0,34 | 0,56 | 0.41 | [0.34-0.5] | |
