## Supplementary Table 4 for "The tree that hides the forest: identification of common predisposing loci in several hematopoietic cancers and several dog breeds"

Supplementary table 4: Association analysis of haplotype of CFA14: 11Mb region with the phenotype in predisposed breeds. The haplotype was determined by the genotype of the following SNVs (11021670, 11023871, 11026224, 11028349, 11030215, 11033259, 11041572, 11067136, 11070626, 11077346, 11083217, 11092391, 11094795, 11108670). At risk haplotypes in the breeds are represented in bold. CI: confidence interval (Woolf Method).

| | Haplotype frequencies in the 3 breeds | | | Odds Ratio | | |
|---|---|---|---|---|---|---|
| | haplotype | affected | unaffected | Odds Ratio | confidence interval (Woolf Method) | pval |
| BMD | **ATAGGAACCCGCGT** | 0,74 | 0,61 | 1.79 | [1.44-2.22] | 9.08x10-8 |
| | ACCAACGCACGCAT | 0,17 | 0,19 | 0.85 | [0.65-1.1] | 0.21 |
| | GCCAACGTATAAGT | 0,05 | 0,15 | 0.34 | [0.24-0.49] | 1.69x10-9 |
| | ATAGGAACCCGCAT | 0,04 | 0,04 | 0.8 | [0.48-1.33] | 0.38 |
| | others | 0,01 | 0,01 | 0.69 | [0.24-2] | 0.48 |
| Rottweiler | **ATAGGAACCCGCGT** | 0,58 | 0,43 | 1.77 | [0.85-3.69] | 0.12 |
| | ATAGGAACCCGCAT | 0,35 | 0,41 | 0.75 | [0.35-1.59] | 0.45 |
| | GCCAACGTATAAGT | 0,06 | 0,11 | 0.56 | [0.15-2.05] | 0.37 |
| | others | | | 0.29 | [0.03-3.29] | 0.55 |
| FCR | **ACCAACGCACGCAC** | 0,46 | 0,2 | 3.43 | [1.05-11.17] | 0.036 |
| | GCCAACGTATAAGT | 0,23 | 0,53 | 0.26 | [0.08-0.83] | 0.02 |
| | ATAGGAACCCGCAC | 0,23 | 0,23 | 0.99 | [0.29-3.44] | 0.98 |
| | others | 0,08 | 0,03 | 2.42 | [0.21-28.34] | 0.59 |
| all breeds | GCCAACGTATAAGT | 0,06 | 0,16 | 0.34 | [0.24-0.47] | 5.37x10-11 |
| | others | 0,94 | 0,84 | 2.92 | [2.1-4.07] | |
