## Supplementary Table 5 for "The tree that hides the forest: identification of common predisposing loci in several hematopoietic cancers and several dog breeds"

Supplementary table 5: Association analysis of CFA05, CFA11 and CFA14 with the phenotype in predisposed breeds. The number of risk alleles was determined with the genotype of the following SNVs (CFA5:30496048, CFA11:41252822, CFA14:11021670) CI: confidence interval (Woolf Method).

| | frequencies of risk alleles in the 3 breeds | | | Odds Ratio | | |
|---|---|---|---|---|---|---|
| | number of risk alleles | affected | unaffected | Odds Ratio | interval (Wo | pval |
| BMD | ≥5 risk alleles | 0,72 | 0,31 | 5.81 | [4.26-7.92] | 1.33x10-30 |
| | 4 risk alleles | 0,23 | 0,4 | 0.45 | [0.33-0.61] | 3.99x10-7 |
| | ≤ 3 risk alleles | 0,05 | 0,29 | 0.13 | [0.08-0.21] | 7.45x10--20 |
| Rottweiler | ≥5 risk alleles | 0,74 | 0,43 | 3.77 | [1.26-11.26] | 0.015 |
| | 4 risk alleles | 0,18 | 0,26 | 0.62 | [0.18-2.14] | 0.45 |
| | ≤3 risk alleles | 0,08 | 0,3 | 0.27 | [0.06-1.15] | 0.03 |
| FCR | 4 risk alleles | 0,38 | 0 | NA | NA | 0.013 |
| | 3 risk alleles | 0,54 | 0,53 | 1.02 | [0.23-4.52] | 1 |
| | ≤2risk alleles | 0,08 | 0,47 | 0.1 | [0.01-0.98] | 0.037 |
| all breeds | ≥5 risk alleles | 0,7 | 0,27 | 5.41 | [4.04-7.24] | 1.14x10-31 |
| | 4 risk alleles | 0,23 | 0,34 | 0.5 | [0.37-0.67] | 2.74x10-6 |
| | ≤3 risk alleles | 0,07 | 0,29 | 0.15 | [0.1-0.23] | 2.17x10-21 |
