## Supplementary Table 6 for "The tree that hides the forest: identification of common predisposing loci in several hematopoietic cancers and several dog breeds"

Supplementary Table 6: Top 10 variants identified in chromosome 11, 14 and 5 loci after imputation of captured variants on 455 HS cases and 408 controls from BMD, Rottweiler and FCR breeds. For chromosome 14 and 5, the association analyses were performed taking in account the information of CFA11 best SNV (CFA11: 41252822).

| Variant | Chromosome | Position | A1 | A2 | P value | Pc1df |
|---|---|---|---|---|---|---|
| AX-167601774_11_41252822 | 11 | 41252822 | T | C | 1.67E-25 | 9.54E-12 |
| indel_49 | 11 | 41216757 | TT | - | 6.43E-25 | 1.70E-11 |
| AX-168062447_11_41215628 | 11 | 41215628 | C | G | 7.39E-25 | 1.81E-11 |
| SNPcapture_11_41175498 | 11 | 41175498 | T | A | 1.23E-24 | 2.26E-11 |
| AX-167216997_BICF2P122855_11_41176819 | 11 | 41176819 | A | G | 1.23E-24 | 2.26E-11 |
| SNPcapture_11_41195083 | 11 | 41195083 | T | G | 1.29E-24 | 2.30E-11 |
| AX-167664066_11_41161357 | 11 | 41161357 | T | G | 1.49E-24 | 2.45E-11 |
| SNPcapture_11_41161357bis | 11 | 41161357 | T | G | 1.49E-24 | 2.45E-11 |
| AX-168281734_BICF2S23134137_11_41161441 | 11 | 41161441 | T | C | 1.49E-24 | 2.45E-11 |
| AX-168107901_11_41163558 | 11 | 41163558 | T | C | 1.49E-24 | 2.45E-11 |
| indel_27623 | 14 | 10951501 | T | - | 1.63E-12 | 6.07E-07 |
| AX-168073054_14_11021670 | 14 | 11021670 | A | G | 1.68E-12 | 6.16E-07 |
| AX-168233524_BICF2P899346_14_11004636 | 14 | 11004636 | A | C | 1.98E-12 | 6.70E-07 |
| SNPcapture_14_11004636 | 14 | 11004636 | A | C | 1.98E-12 | 6.70E-07 |
| AX-167740817_14_11004924 | 14 | 11004924 | G | T | 1.98E-12 | 6.70E-07 |
| SNPcapture_14_11025603 | 14 | 11025603 | A | G | 1.98E-12 | 6.70E-07 |
| indel_27704 | 14 | 11027076 | T | - | 1.98E-12 | 6.70E-07 |
| indel_27716 | 14 | 11035998 | A | - | 1.98E-12 | 6.70E-07 |
| SNPcapture_14_11043035 | 14 | 11043035 | G | A | 1.98E-12 | 6.70E-07 |
| AX-167368660_14_11067136 | 14 | 11067136 | C | T | 1.98E-12 | 6.70E-07 |
| SNPcapture_5_30488886 | 5 | 30488886 | G | A | 1.63E-11 | 1.96E-06 |
| SNPcapture_5_30489203 | 5 | 30489203 | G | C | 2.02E-11 | 2.18E-06 |
| SNPcapture_5_30489217 | 5 | 30489217 | T | C | 2.02E-11 | 2.18E-06 |
| AX-167250226_BICF2G630182217_5_30496048 | 5 | 30496048 | T | C | 2.39E-11 | 2.38E-06 |
| SNPcapture_5_30491343 | 5 | 30491343 | C | T | 2.42E-11 | 2.39E-06 |
| AX-167199530_BICF2G630182204_5_30488801 | 5 | 30488801 | T | A | 2.46E-11 | 2.41E-06 |
| SNPcapture_5_30483338 | 5 | 30483338 | G | A | 2.52E-11 | 2.44E-06 |
| SNPcapture_5_30485505 | 5 | 30485505 | A | G | 2.52E-11 | 2.44E-06 |
| SNPcapture_5_30486583 | 5 | 30486583 | G | A | 2.52E-11 | 2.44E-06 |
| SNPcapture_5_30488559 | 5 | 30488559 | G | A | 2.52E-11 | 2.44E-06 |
