## Supplementary Table 7 for "The tree that hides the forest: identification of common predisposing loci in several hematopoietic cancers and several dog breeds"

Supplementary Table 7:List of candidate genes identified by the GWAS performed on several dog breed and several cancers. The gene indicated corresponds to the closest genes to the best SNV per GWAS experiment. The involvement of these genes in human cancer or inflammation is summarized according to NIH gene database ( https://www.ncbi.nlm.nih.gov/gene); the association to specific traits in human GWAS is summarized according to phegeni database, (https://www.ncbi.nlm.nih.gov/gap/phegeni).

| Gene | breeds | cancer | GWAS | involved in human cancer | | involved in inflammation | pathway | Trait association |
|---|---|---|---|---|---|---|---|---|
| | | | | predisposition | somatic disregulation | | | |
| SPNS3 | BMD<br>FCR | HS<br>lymphoma | GWAS_1_HS_BMD<br>GWAS_2_HS+lymphoma_BMD<br>GWAS_8_HS_BMD_with_imputed_SNV<br>GWAS_9_HS_BMD+FCR_with_imputed_SNV | yes (glioma, melanoma, chronic lymphocytic leukemia) | yes (chronic lymphocytic leukaemia, breast cancers) | | Cell Cycle, DNA Damage/Telomere Stress Induced Senescence, Chromosome Maintenance | Chemokine CCL2, Platelet Function Tests, Erythrocyte Count,Eosinophils |
| MTAP | BMD<br>Rottweiler | HS | GWAS_1_HS_BMD<br>GWAS_7_HS_BMD+Rottweiler | yes (melanoma, Diaphyseal medullary stenosis with malignant fibrous histiocytoma) | yes (deficient in many cancers because this gene and the tumor suppressor p16 gene are co-deleted) | yes | Interleukin-12 signaling,Cytokine Signaling in Immune system | Blood Pressure, Melanoma, Monocytes, cholesterol, LDL |
| FHIT | BMD | HS<br>mast cell tumor | GWAS_1_HS_BMD<br>GWAS_4_HS+MCT_BMD | yes (breast cancer) | yes ( in numerous cancer : breast cancer, bladder cancer…) | | | Blood Pressure, Cholesterol, LDL,Fibrinogen, Platelet Function Tests, Platelet Function Tests, Precursor Cell Lymphoblastic Leukemia-Lymphoma |
| TRPC6 | BMD<br>golden retriever | HS<br>lymphoma | GWAS_3_HS+lymphoma_BMD+golden_retriever | | | | | Hematocrit, Platelet Function Tests |
| BORCS6 | BMD<br>golden retriever | HS<br>lymphoma | GWAS_3_HS+lymphoma_BMD+golden_retriever | | | | mTORC1 | Breast Neoplasms |
| ARHGEF3 | BMD | HS<br>mast cell tumor | GWAS_4_HS+MCT_BMD | | yes (in acute myeloid leukemias, carcinoma) | | mTORC2 | Mean Platelet Volume, Platelet Count,Platelet Function Tests, Arthritis, Rheumatoid |
| C3orf67 | BMD<br>golden retriever | HS<br>mast cell tumor | GWAS_5_HS+MCT_BMD+golden_retriever | | | | | |
| IL17RD | BMD<br>golden retriever | HS<br>mast cell tumor | GWAS_5_HS+MCT_BMD+golden_retriever | | yes (in colorectal cancer, regulation of Epithelial-Mesenchymal Transition in Breast Cancer or prostate cancer, ) | yes (it is an inhibitor of proinflammatory cytokine signaling, acting by cytoplasmic sequestration of NF-κB) | Immune system, MAPK kinase activation,IL17 signaling pathway | Platelet Function Tests |
| BRSK2 | BMD | HS<br>mast cell tumor | GWAS_4_HS+MCT_BMD | | yes (in pancreatic cancer) | | mTORC1 | Albumins |
| CAAP1 | BMD<br>FCR | HS | GWAS_6_HS_BMD+FCR | | | | Apoptosis | Asthma, Allergens, Platelet Function Tests, Gallbladder Neoplasms,Triglycerides,Hematocrit,Subcutaneous Fat, Prostatic Neoplasms, Cholesterol |
| CDKN2A | BMD<br>Rottweiler | HS | GWAS_7_HS_BMD+Rottweiler<br>GWAS_10_HS_BMD+FCR+Rottweiler_with_imputed_SNV | yes (melanoma, astrocytoma pancreatic cancer) | yes (This gene is frequently mutated or deleted in a wide variety of tumors, and is known to be an important tumor suppressor gene) | | Apoptosis, cell cycle, | Platelet Count, Precursor B-Cell Lymphoblastic Leukemia-Lymphoma,Gastrointestinal Neoplasms, Glioma, Basophils |
| CDKN2B | BMD<br>Rottweiler | HS | GWAS_7_HS_BMD+Rottweiler<br>GWAS_10_HS_BMD+FCR+Rottweiler_with_imputed_SNV | | yes (ovarian cancer, T-acute lymphoblastic leukemia) | | cell cycle, signaling TGF-beta | Glioma, Gastrointestinal Neoplasms |
| CDKN2B_AS1 | BMD<br>Rottweiler | HS | GWAS_7_HS_BMD+Rottweiler<br>GWAS_10_HS_BMD+FCR+Rottweiler_with_imputed_SNV | | yes (liver cancer) | associated with Inflammatory Bowel Disease and participates in intestinal barrier | | Glioma, Carcinoma, Basal Cell,Mouth Neoplasms,Breast Neoplasms, Prostatic Neoplasms,Lung Neoplasms,Ovarian Neoplasms,Prostatic Neoplasms, Nasopharyngeal Neoplasms,Esophageal Squamous Cell Carcinoma, Gastrointestinal Neoplasms |
| PSMD4 | BMD<br>FCR | HS | GWAS_6_HS_BMD+FCR | | yes (hepatocellular carcinoma.) | subunit of proteasome ibvolved in processing of class I MHC peptides | APC/C-mediated degradation of cell cycle proteins,DNA Replication, Adaptive Immune System, Activation of NF-kappaB in B cells, C-type lectin receptors (CLRs), Cytokine Signaling in Immune system, Downstream TCR signaling, Downstream signaling events of B Cell Receptor (BCR), regulation of Apoptosis | |
| PFKFB3 | BMD<br>FCR<br>Rottweiler | HS | GWAS_10_HS_BMD+FCR+Rottweiler_with_imputed_SNV | | yes (colon cancer ,non-small cell lung cancer, breast cancer) | yes regulation of macrophage, critical role in TNF-alpha-induced endothelial inflammation. | cell cycle, apoptosis | Breast Neoplasms,Hypertension, Ferritins,Cholesterol, HDL |
| MPDZ | BMD<br>FCR<br>Rottweiler | HS | GWAS_10_HS_BMD+FCR+Rottweiler_with_imputed_SNV | | | | | Cholesterol, LDL |
| IL17A | BMD<br>FCR | HS | GWAS_9_HS_BMD+FCR_with_imputed_SNV | | yes (hepatocellular carcinoma, gastric cancer.nasopharyngeal carcinoma..) | yes High levels of this cytokine are associated with several chronic inflammatory diseases including rheumatoid arthritis, psoriasis and multiple sclerosis | Immune system, IL17 signaling pathway | Hematocrit,Platelet Function Tests, Hemoglobins |
| LEP | BMD | HS | GWAS_8_HS_BMD_with_imputed_SNV | | | yes involved in the regulation of immune and inflammatory responses, hematopoiesis | | Platelet Function Tests, Erythrocyte Count, Blood Pressure |
| CPA1 | BMD<br>FCR | HS | GWAS_9_HS_BMD+FCR_with_imputed_SNV | yes (in panceratic cancer) | yes (in panceratic cancer) | yes involved in chronic pancreatitis | | |
| POT1 | BMD<br>FCR<br>Rottweiler | HS | GWAS_10_HS_BMD+FCR+Rottweiler_with_imputed_SNV | yes (Hodgkin lymphoma, melanoma, glioma, chronic lymphocytic leukemia) | yes( in Chronic lymphocytic leukemia ) | | Cell Cycle, Chromosome Maintenance, Cellular Senescence, DNA Repair,Telomere Maintenance | Leukemia, Lymphocytic, Chronic, B-Cell, Platelet Function Tests, |
| DDX55 | BMD | HS | GWAS_8_HS_BMD_with_imputed_SNV | | | | | |
| TBL1X | BMD | HS | GWAS_8_HS_BMD_with_imputed_SNV | yes (in prostate cancer) | yes (in prostate cancer) | yes, inhibits NF-κB-target gene expression and suppresses inflammatory signaling | Chromatin organization, Infectious disease, Metabolism of lipids | Cholesterol, HDL |
| SHROOM2 | BMD | HS | GWAS_8_HS_BMD_with_imputed_SNV | yes (colorectal cancer and prostate cancer ) | | | | Prostatic Neoplasms,Colorectal Neoplasms |
| TUSC1 | BMD<br>FCR<br>Rottweiler | HS | GWAS_10_HS_BMD+FCR+Rottweiler_with_imputed_SNV | | yes( in non-small-cell lung cancer and small-cell lung cancer , glioblastoma, gastric cancer, hepatocellular carcinoma) | | | Blood Circulation, Rhinitis, Allergic, Platelet Function Tests,Hematocrit, Erythrocyte Count,Blood Pressure,Erythrocyte Indices, Interleukin-10, Hematocrit, Cholesterol, HDL, |
| C9orf72 | BMD<br>FCR<br>Rottweiler | HS | §3.5 | | | yes, Null mice developed a robust immune phenotype characterized by myeloid expansion, T cell activation, and increased plasma cells | | |
